# Quartet RNA reference materials and ratio-based reference datasets for reliable transcriptomic profiling

**DOI:** 10.1101/2022.09.26.507265

**Authors:** Ying Yu, Wanwan Hou, Haiyan Wang, Lianhua Dong, Yaqing Liu, Shanyue Sun, Jingcheng Yang, Zehui Cao, Peipei Zhang, Yi Zi, Zhihui Li, Ruimei Liu, Jian Gao, Qingwang Chen, Naixin Zhang, Jingjing Li, Luyao Ren, He Jiang, Jun Shang, Sibo Zhu, Xiaolin Wang, Tao Qing, Ding Bao, Bingying Li, Bin Li, Chen Suo, Yan Pi, The Quartet Project Team, Xia Wang, Fangping Dai, Andreas Scherer, Pirkko Mattila, Jingxiong Han, Lijun Zhang, Hui Jiang, Danielle Thierry-Mieg, Jean Thierry-Mieg, Wenming Xiao, Huixiao Hong, Weida Tong, Jing Wang, Jinming Li, Xiang Fang, Li Jin, Leming Shi, Joshua Xu, Feng Qian, Rui Zhang, Yuanting Zheng

**Author notes:** Corresponding authors’ e-mail addresses. These authors contributed equally: Ying Yu, Wanwan Hou, Haiyan Wang, Lianhua Dong, Yaqing Liu.

## Abstract

As an indispensable tool for transcriptome-wide analysis of differential gene expression, RNA sequencing (RNAseq) has demonstrated great potential in clinical applications. However, the lack of multi-group RNA reference materials of biological relevance and the corresponding reference datasets for assessing the reliability of RNAseq hampers its wide clinical applications wherein the underlying biological differences among study groups are often small. As part of the Quartet Project for quality control and data integration of multiomic profiling, we established four RNA reference materials derived from immortalized B-lymphoblastoid cell lines from four members of a monozygotic twin family. Additionally, we constructed ratio-based transcriptome-wide reference datasets using multi-batch RNAseq datasets, providing “ground truth” for benchmarking. Moreover, Quartet-sample-based quality metrics were developed for assessing reliability of RNAseq technology in terms of intra-batch proficiency and cross-batch reproducibility. The small intrinsic biological differences among the Quartet samples enable sensitive assessment of performance of transcriptomic measurements. The Quartet RNA reference materials combined with the reference datasets can be served as unique resources for assessing data quality and improving reliability of transcriptomic profiling.

---

RNA sequencing (RNAseq) is an indispensable tool for transcriptome-wide analysis of differential gene expression and is widely used in biomedical research to discover biomarkers for clinical diagnosis, prognosis, and therapeutic action^1–8^. As RNAseq-based biomarker discovery continues to advance, RNAseq-based routine clinical utility is expected to become a reality^4, 9^. For example, clinical tests complemented by measuring the differential expression of clinically relevant genes will facilitate the prediction of clinical outcomes and treatment decisions^5, 10, 11^. It should be noticed that clinically relevant differences in gene expression among study groups are often small^6^. Hence, there is a consistent need for making RNAseq more reliable to enhance its power of detecting differential expression, especially for clinical applications. The reliability of RNAseq technology comprises two aspects. First, intra-batch (or lab) proficiency is a prerequisite for ensuring that data from a certain lab or batch are acquired with the best proficiency obtainable with the technology^12^. Secondly, cross-batch reproducibility is required for ensuring closely similar differential expression results from replicate samples processed with different platforms, labs, protocols or batches^13^. Cross-batch reproducibility also refers to multi-batch integrability that means the ability to provide closely similar results between within-batch analysis and cross-batch integrative analysis in the existence of widespread batch effects^14, 15^.

Reference materials are valuable tools for evaluating the reliability of omic data^16, 17^. Based on RNAseq data generated with reference materials from different platforms, labs or batches, reliability can be objectively evaluated according to the two aforementioned aspects. The MicroArray/Sequencing Quality Control (MAQC/SEQC) consortia have established two publicly available transcriptome-wide RNA reference materials that are derived from ten cancer cell lines (sample A, Universal Human Reference RNA) and brain tissues of 23 donors (sample B, Human Brain Reference RNA)^13^. Based on these RNA reference materials, the MAQC/SEQC consortia systematically evaluated the performances of different platforms and labs in utilizing the microarray^13^ and RNAseq^18^ technologies. The results revealed generally good intra-platform reproducibility across test sites as well as high cross-platform concordance, according to differentially expressed genes and their fold-changes of expression levels between the two RNA samples^13, 18, 19^. Furthermore, the MAQC RNA reference materials have been served as resources for the research community to develop and validate new RNA quantification technologies^20^.

Despite of the successful applications of the MAQC RNA reference materials, several challenges remain. First, the huge biological differences between the two MAQC reference materials^21^ are substantially larger than groupwise differences commonly seen in most clinically relevant scenarios. Distinguishing the two MAQC RNA reference materials is easy and straightforward, but does not automatically guarantee the performance needed for differentiating more subtle differential expression in clinical scenarios. Secondly, the ability of distinguishing two sample groups does not translate to the ability of reliably distinguishing more than two sample groups as commonly seen in clinical applications. Thirdly, the current stock of the MAQC B sample is almost exhausted, and it is difficult to be regenerated. Therefore, there is an urgent need for multiple RNA reference materials with subtle inter-sample differences, high stability, long-term availability, and easy manufacturability.

Furthermore, reference datasets can be used as “ground truth” in performance assessment. Prior studies have shown that genome-wide reference datasets of genetic variants enable improvement of the reproducibility and accuracy of clinical applications of cancer^22, 23^ and genetic diseases^24, 25^. However, there is a paucity of transcriptome-wide reference datasets^4, 16^. Therefore, transcriptome-wide reference datasets associated with publicly available RNA reference materials are urgently needed but are lacking^16^.

As a part of the Quartet Project for the quality control and data integration of multiomic profiling (http://chinese-quartet.org/), we established four RNA reference materials derived from immortalized cell lines from the four members of a monozygotic twin family quartet, which exhibited subtle inter-sample differences, high stability, long-term availability, and easy manufacturability. Furthermore, matched multiomic reference materials including DNAs^26^, proteins^27^ and metabolites^28^ were established along with RNAs from the same culturing of the immortalized cell lines (see accompanying manuscripts elsewhere). The comprehensive set of multiomic reference materials enables cross-omics validations and performance assessment at multiomic integration level^29^. Moreover, we performed a multi-lab RNAseq study comprising 21 batches of data. Based on the high-quality multi-lab RNAseq datasets, we established the ratio-based reference datasets of gene expression via consensus, and developed quality metrics for assessing reliability of RNAseq technology in terms of intra-batch proficiency and cross-batch reproducibility. The Quartet RNA reference materials and reference datasets along with the quality metrics provided in this study can be served as unique resources for lab proficiency testing, objectively assessing data quality of transcriptomic profiling studies, and improving the reliability of cross-batch integrative analysis.

## Results

### Overview of study design

The Quartet RNA reference materials were derived from the Epstein-Barr Virus (EBV) immortalized B-lymphoblastoid cell lines from four members of a Chinese family quartet including monozygotic twin daughters (D5 and D6), father (F7), and mother (M8) (Fig. 1a). Large quantities of RNA (over five milligrams) were obtained per cell line, enabling standard RNAseq experiments over 10,000 to 50,000 times and providing a material basis for long-term quality monitoring. RNA quality was high according to RNA integrity number (RIN) and RNA purity (Extended Data Fig. 1 and **Supplementary Table 1)**. Moreover, the RNA reference materials showed adequate stability across 20 months of storage at −80°C, or 14 days of storage at room temperature (25 °C) or 4°C, or up to 20 times of bottle-opening and freeze-thaw cycle (Extended Data Fig. 2).

**Figure 1.**
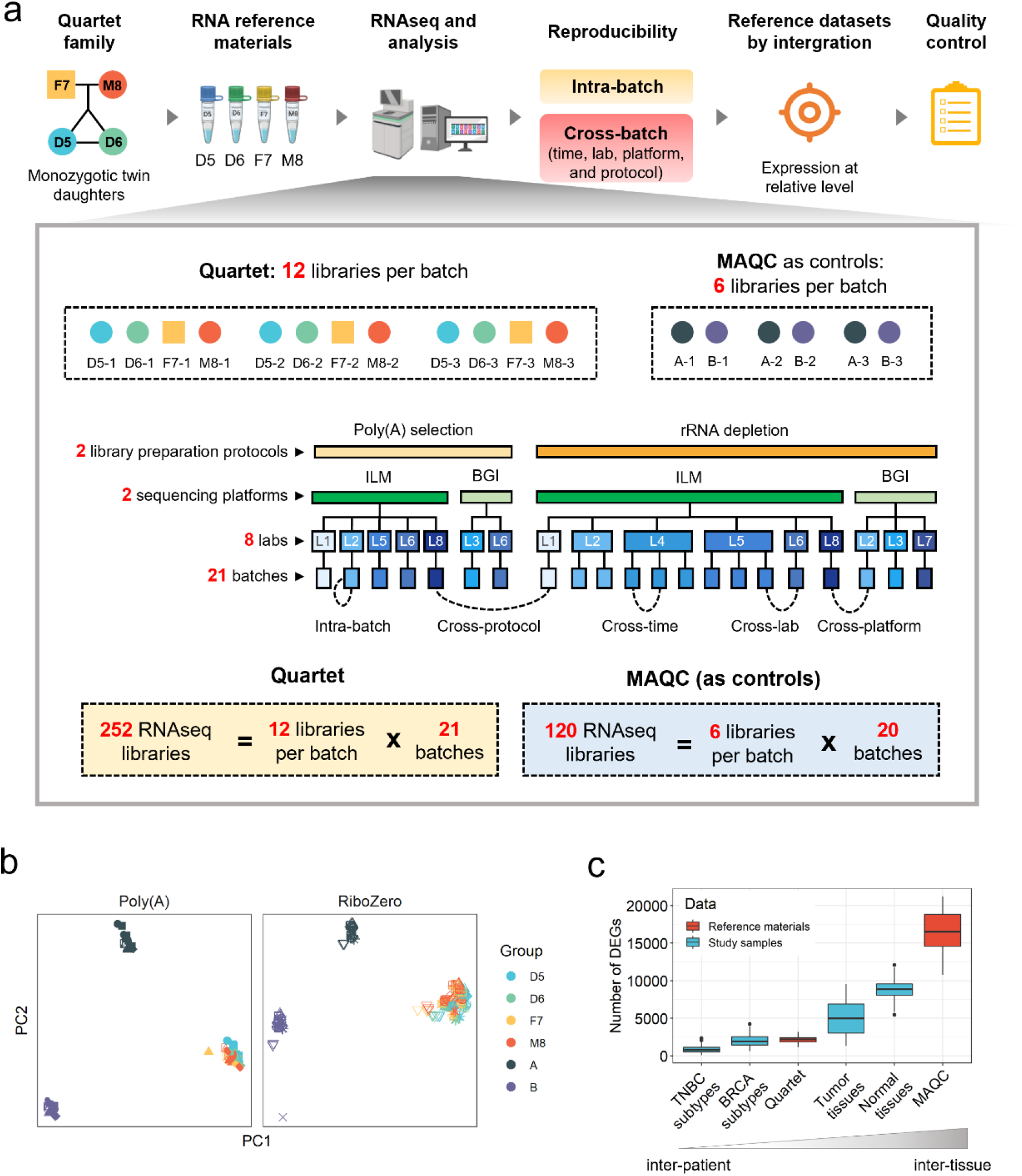
Overview of study design. (**a**) Quartet RNA reference materials were derived from immortalized Epstein-Barr Virus (EBV) infected B-lymphoblastoid cell lines from a quartet family including monozygotic twin daughters (D5 and D6), and their father (F7) and mother (M8). Multi-batches of RNAseq datasets were generated from independent labs using different library preparation protocols and sequencing platforms. Intra-batch proficiency and cross-batch reproducibility were then estimated. Based on multi-batches of RNAseq data, we constructed ratio-based transcriptome-wide reference datasets, and developed corresponding quality metrics. (**b**) Scatterplots of principal components (PCs) on RNAseq data of the Quartet and MAQC RNA reference materials (marked in colors) in the 21 batches (marked in shapes). (**c**) Boxplots showing the numbers of differentially expressed genes (DEGs) among Quartet reference materials, MAQC reference materials, and four clinical/biological classification problems from published data sets. The four clinical/biological classifications used to represent clinical scenarios include four subtypes of triple-negative breast cancers (TNBC) with different therapeutic actions^7^, four subtypes of breast cancers (BRCA) with different prognosis and therapeutic actions (luminal A, luminal B, basal-like/triple negative, and HER2-positive)^30^, four types of tumor tissues^30^, and four types of normal tissues^31^. The latter two types of biological classification problems are important for understanding the genetic basis of human diseases. A gene was identified as differentially expressed when satisfying the criteria of Student’s t-test *p* < 0.05 and fold change ≥ 2 or ≤ 0.5 between two groups or conditions.

RNAseq datasets from the Quartet RNA reference materials were then obtained, consisting of 252 RNAseq libraries from 21 batches generated in eight labs using two library construction protocols (poly(A) selection and RiboZero) and two sequencing platforms (Illumina NovaSeq and BGI DNBseq) (Fig. 1a and **Supplementary Table 2**). Here, a batch is defined as 12 libraries from a standard sample set, consisting of 12 tubes with each representing one of the triplicates of the Quartet RNA reference sample groups, whose library construction and sequencing experiments were conducted simultaneously. On the other hand, libraries conducted at different time points, in different labs, with different sequencing platforms, or using different library preparation protocols are recognized broadly as cross-batch libraries (Fig. 1a). This comprehensive study design allows for objective performance assessment at multiple levels including cross-time, cross-lab, cross-platform, and cross-protocol. Moreover, RNAseq experiments with the MAQC RNA reference materials (A and B) were conducted simultaneously with the Quartet reference materials in 20 of the 21 batches (Fig. 1a), enabling head-to-head comparisons between the two sources (MAQC versus Quartet) of RNA reference materials. In addition, bioinformatic analysis pipelines were validated using data from the MAQC RNA reference materials by comparison with previous researches^13, 18^ (Extended Data Fig. 3).

### Quartet RNA reference materials exhibit small intrinsic biological differences

Using principal component analysis (PCA) as an exploratory overview of data analysis, we found that multi-batch libraries of the Quartet reference materials from the same protocol (poly(A) or RiboZero) were clustered together, whereas libraries of MAQC A and B samples were clustered separately into two distinct groups (Fig. 1b). This result indicates that the intrinsic biological differences among the four groups of Quartet RNA reference materials are much smaller compared to those between the two MAQC RNA reference materials.

To investigate whether the magnitude of intrinsic biological differences or signals between the Quartet reference materials is representative of those seen in clinically relevant scenarios, we compared the extent of intrinsic biological differences between reference materials (MAQC A versus B, and Quartet members) and those of four biological classification problems from published datasets ranging from four subtypes of triple-negative breast cancers^7^, four subtypes of breast cancers (luminal A, luminal B, basal-like/triple negative, and HER2-positive)^30^, four types of tumor tissues^30^, and four types of normal tissues^31^. The number of differentially expressed genes (DEGs), previously used as a measure of “treatment effect size”^6^, identified from the four biological classification problems ranged from 884 to 4,980 (mean), corresponding to an increase of intrinsic biological differences and/or decrease of within-group heterogeneity (Fig. 1c). Importantly, the differences among Quartet RNA reference materials were 2,164 ± 424 (mean ± SD) in terms of DEGs, which were ranked in the middle of these four clinical classification scenarios. In contrast, the differences between the two MAQC RNA reference materials were much larger (16,503 ± 2,831, mean ± SD) than the aforementioned biological classification problems (Fig. 1c). These data again illustrate that the intrinsic biological differences among the Quartet reference materials are much smaller than those between MAQC RNA reference materials A and B, and such small differences are comparable to those seen in clinical and biological classification scenarios.

### Signal-to-Noise Ratio (SNR) enables assessment and diagnosis of data quality

Based on the Quartet design, a Signal-to-Noise Ratio (SNR) metric was established to gauge the performance of a platform, a lab, a protocol, or a batch in distinguishing the intrinsic biological differences (“signal”) among the Quartet samples from variations among technical replicates of the same sample group (“noise”) (Fig. 2a). Generally, a lower SNR value indicates lower discriminating power, *vice versa*. For an SNR value around or below zero, it means that the magnitude of signal is at a similar level as the noise or even lower than the noise. In this case, it is almost impossible to distinguish different sample groups under the high level of technical noises (Fig. 2b).

**Figure 2.**
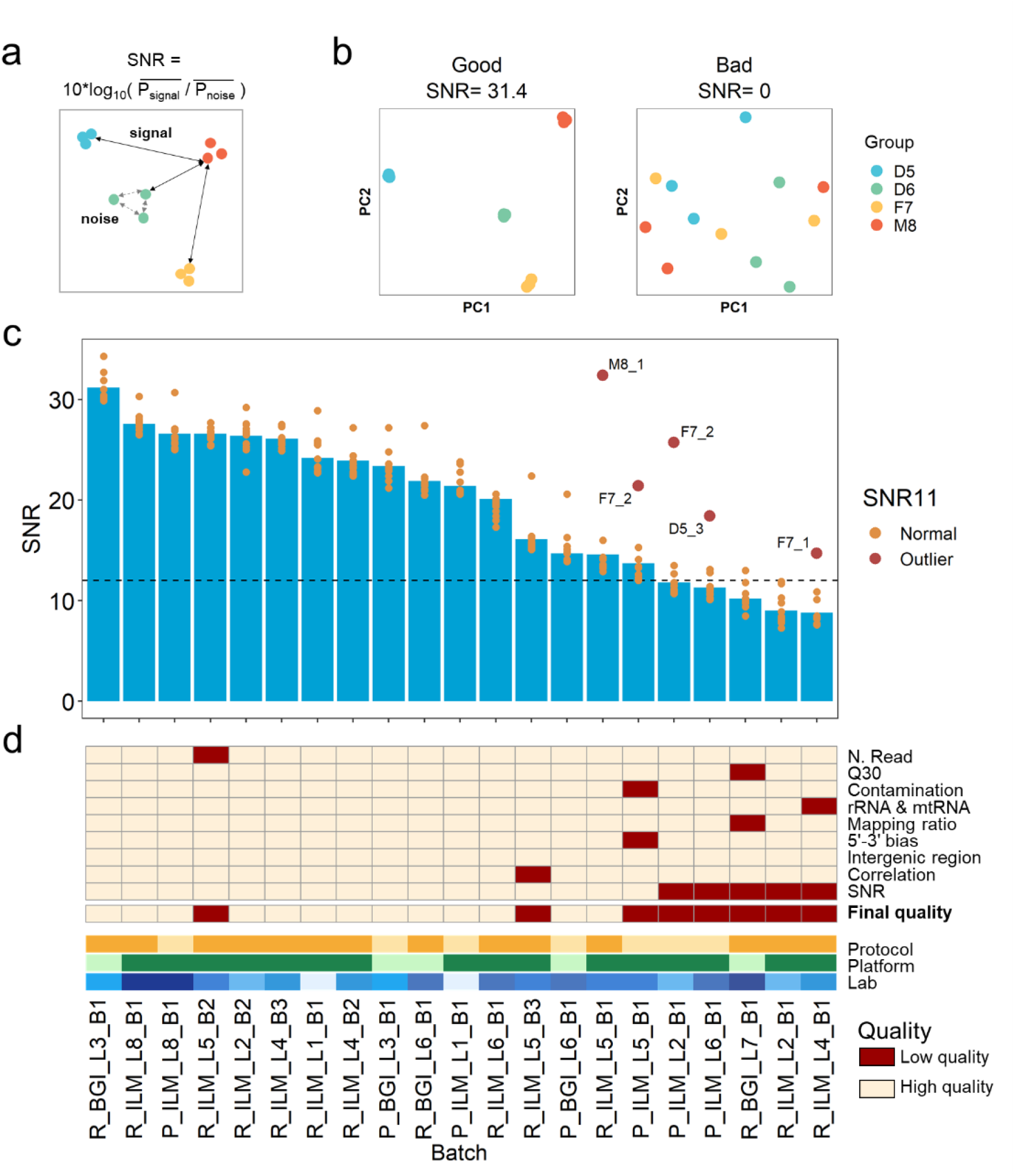
Signal-to-noise ratio enables assessment and diagnosis of data quality. (**a**) Concept of calculating signal-to-noise ratio (SNR). SNR was established to characterize the ability of a platform, a lab, or a batch to distinguish the intrinsic differences among distinct biological sample groups (“signal”) from variations in technical replicates of the same sample group (“noise”). (**b**) Examples of good and bad batches with their SNR values and corresponding principal components analysis (PCA) scatter plots. (**c**) SNR values across 21 RNAseq batches to measure data quality. Batches were ordered by SNR values. Dots represented SNR values based on any 11 of the 12 libraries in each batch. A dot in dark red represented SNR11 value that increased over five decibels compared to its standard SNR (12-sample SNR), when one library in this batch was excluded (the library ID was labeled), while a dot in orange represented SNR11 value that decreased or increased less than five decibels compared to its standard SNR. (**d**) Quality flags of RNAseq batches in terms of the number of sequencing reads (N. Read), percentage of Q30 (Q30), percentage of reads that were mapped to contamination species (e.g. virus, bacteria, fungi) (contamination), percentage reads that were mapped to rRNA or mtRNA (rRNA & mtRNA), percentage of reads that were mapped to the human genome (mapping ratio), Gene body (5’-3’) bias (5’-3’ bias), percentage of mapped reads that were located in intergenic region in human genome (intergenic region), Pearson correlation coefficient of technical replicates (correlation), signal-to-noise ratio (SNR), and final quality flag. Batches were ordered by SNR values. Batch information of the datasets was shown by the color legend.

We evaluated the performance of five different methods in characterizing SNR, including Euclidean distance (Dist), overall expression profiles (Expr), Pearson correlation coefficient (Cor), t-Distributed Stochastic Neighbor Embedding (tSNE), and PCA. It was found that PCA-based SNR outperformed the other four methods in terms of its sensitivity in differentiating the quality of different datasets (Extended Data Fig. 4a). We next computed SNR using different numbers of principal components of PCA (Extended Data Fig. 4b). SNR values based on the first component, first two components, or first three components were highly correlated. Therefore, we used the first two principal components for calculating SNR, in correspondence with visualization in commonly used two-dimensional PCA plots (Figs. 2a and 2b).

SNR enables assessment of quality across the 21 batches of RNAseq data. For most batches, the three replicates from the same sample group can be clearly distinguished from those of other sample groups (Extended Data Fig. 5). Large fluctuations of SNR values were observed across batches generated with the same protocol, same sequencing platform, or even from the same lab, highlighting the needs for objectively assessing and monitoring the technical competency in data generation.

SNR can also be applied to diagnose potential causes of quality issues. In addition to the SNR values considering all 12 libraries in a batch, we also calculated SNR11 values with any 11 of the 12 libraries in each batch (Fig. 2c). In two of the five aforementioned batches with low quality (L2_B1 and L6_B1), the SNR11 values increased by more than 5 decibels compared to the 12-sample SNR values, indicating that the lower SNR values from these two batches might be a result of a “random failure” of a particular technical replicate, e.g., replicate F7-2 from batch L2_B1 and replicate D5-3 from batch L6_B1. In contrast, the other three low-quality batches were possibly due to systematic technical issues, because excluding any specific replicate (or potential outlier) could not significantly improve the SNR values.

Moreover, SNR enables assessment of data quality not only at gene expression level, but also at alternative splicing (AS) level. Similarly, SNR values at AS level varied significantly across batches. SNR values could be as high as 32.3 so that the three technical replicates for each sample type on the PCA plot could be loosely regarded as one dot (Extended Data Fig. 6a), or as low as 2.4 where technical replicates were mixed with libraries from other sample types (Extended Data Fig. 6b).

Using multiple metrics, including SNR and other widely-used quality metrics with fastq, bam, and expression profiles (**Supplementary Table 3**), with SNR showing the greatest differentiating power, 13 batches were flagged as high quality and were used for subsequent data integration to create the reference datasets, whereas the other eight batches were flagged as low quality and excluded from creating reference datasets (Fig. 2d and **Supplementary Table 3**).

### Construction and validation of ratio-based reference datasets

We next constructed transcriptome-wide reference datasets based on multi-batch and high-quality RNAseq datasets, providing “ground truth” for benchmarking. Ratio-based expression profiles, defined as a fold-change or a fold-difference of expression levels between two sample groups for the same gene, agreed well across multiple transcriptomic technologies, including RNAseq, microarray, and qPCR^13, 18^. On the other hand, the incomparability of conventional “absolute” expression profiles across different batches prevented meaningful cross-batch data integration^13, 18^. Hence, we constructed the ratio-based transcriptome-wide reference datasets (Fig. 3a).

**Figure 3.**
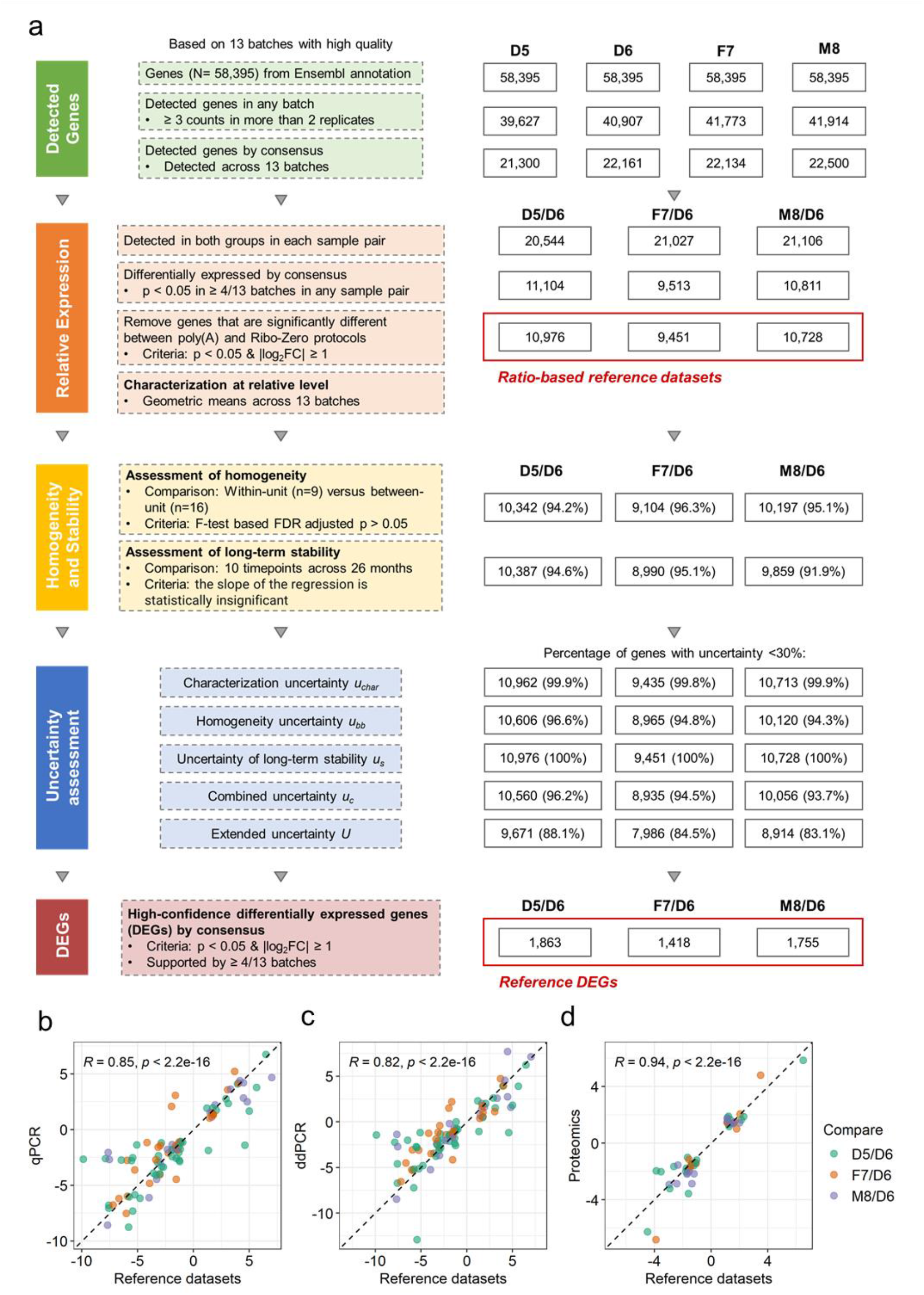
Construction and validation of ratio-based transcriptome-wide reference datasets. (**a**) Workflow for constructing Quartet RNA reference datasets. Reference datasets were constructed according to the following steps: (1) identifying detectable genes; (2) calculating ratio-based expression based on reliably detectable genes that were differentially expressed and with low uncertainty; (3) assessing the homogeneity and stability of RNA reference materials; and (4) assessing the uncertainty of ratio-based reference datasets; (5) identifying high-confidence differentially expressed genes (DEGs) in reference datasets. (**b-d**) Scatter plots of log2 fold changes (FC) of gene expression between reference datasets and qPCR (**b**), ddPCR (**c**) and proteomics data (**d**). Pearson correlation coefficient across three sample pairs was calculated. Genes that were considered as DEGs in both methods shown in X-axis and Y-axis were used for plotting. X-axis: average log2FC from 13 high-quality RNAseq batches of high-confident DEGs from reference datasets. Y-axis: for the qPCR and proteomics data, a gene or a protein was considered as a DEG when the t-test *p* < 0.05 and fold change ≥ 2 or ≤ 0.5; for ddPCR data, genes that were identified as DEGs based on qPCR were used. Average log2FC of qPCR (n=3), ddPCR (n=2), and proteomics (n=3) data from DEGs were used for plotting.

First, the detectable genes in each sample group (D5, D6, F7, and M8) were identified by consensus separately. Briefly, if a gene was detected in all of 13 batches in a sample group, it was considered expressed in that sample group. For the four Quartet reference materials (D5, D6, F7, and M8), 21,300, 22,161, 22,134, and 22,500 genes were expressed, respectively, representing 36.5% to 38.5% of the 58,395 genes annotated in GRCh38 (Fig. 3a).

Secondly, ratio-based expressions (as log2 transformed) were calculated for three pairs of sample groups using replicates of D6 as the common denominator (D5/D6, F7/D6, and M8/D6). In order to improve the reliability of the reference values, genes that were satisfied with thresholds of *p* < 0.05 in each sample pair were used. Furthermore, genes that were significantly different between poly(A) and RiboZero protocols were removed to minimize technical variations introduced by the differences between the two distinct library preparation protocols. After these filtrations, the number of retained genes were 10,976, 9,451 and 10,728 for the three sample pairs (D5/D6, F7/D6, and M8/D6) (Fig. 3a). Ratio-based reference datasets were then characterized between each pair of samples for a gene and were provided in the format of a geometric mean by summarizing from the 13 fold-changes calculated from each of the 13 high-quality RNAseq datasets (**Supplementary Tables 4** and **5**).

Thirdly, the homogeneity and stability of the reference materials were assessed (Fig. 3a and Extended Data Fig. 7). Homogeneity and stability are two crucial characteristics of reference materials^32^. Homogeneity assessment aims to ensure that the previously characterized properties of reference materials are uniformly distributed across packaging units of the reference materials. Because the Quartet reference materials were characterized using gene expressions, homogeneity assessment was conducted based on gene expression data. Here, we evaluated the homogeneity of reference materials by calculating within-unit (n=9) versus between-unit (n=16) variances of each gene using the ANOVA method (Extended Data Figs. 7a and 7b). Most (94.2%-96.3%) genes performed well in homogeneity assessment (**Supplementary Table 6**). On the other hand, stability assessment aims to ensure that the value of the properties previously characterized remains unchanged over time. Here, we evaluated stability of the Quartet reference materials by calculating ANOVA variances of each gene based on 15 batches of RNAseq datasets that were generated over 26 months (Extended Data Figs. 7c and 7d). Most genes (91.9%-95.1%) performed well in long-term stability assessment (**Supplementary Table 6**). Therefore, the Quartet RNA reference materials stored at −80°C were considered to be homogenous and stable, as can be seen from the corresponding reference datasets.

Fourthly, uncertainties of reference materials were estimated. It is essential for identifying each source of uncertainty and to quantify the uncertainty introduced by each source. According to ISO Guide 35 (2017)^32^ and JJF-1343 (2012)^33^, the source of uncertainties can be classified into characterization uncertainties (*u_char_*), sample inhomogeneities (between-bottle variation, *u_bb_*), and instabilities (*u_s_*). These values were then combined to form the combined uncertainties (*u_c_*) and expanded uncertainties (*U*) with an expansion factor (*k*=2, 95% confidence level) (Fig 3a, and **Supplementary Table 6**). As a result, the majority of genes (83.1%∼ 88.1%) showed a limited expanded uncertainties of less than 30%, demonstrating that the characterization of reference datasets was valid.

Finally, high-confidence DEGs in the reference datasets (reference DEGs) were identified. A gene was considered as a reference DEG between two sample types if it was concordantly discovered as an up- or down-regulated gene (*p* < 0.05 and fold change ≥ 2 or ≤ 0.5) in more than six of the 13 high-quality batches. The number of reference DEGs ranged from 1,418 to 1,863 for the three sample pairs (Figs. 3a and **Supplementary Table 7**).

To verify the reliability of the reference datasets, we performed qPCR as an orthogonal validation. We selected 82 genes from the Quartet reference datasets and conducted qPCR experiments on the four reference materials (**Supplementary Table 8**). There is a high level of concordance between the Quartet reference datasets and the qPCR data in terms of DEGs (92%, 91 of 99 DEGs across three sample pairs). We also compared the fold change of qPCR versus that of reference datasets for the DEGs that were detected by both technologies (n=91). We observed an expected high level of concordance to qPCR (*r=*0.85), similar to what were previously reported (*r*=0.80∼1)^6^ (Fig. 3b and **Supplementary Table 9**). DEGs that were identified in the reference datasets and qRT-PCR were further validated using droplet digital PCR (ddPCR). Similar result was observed when comparing the fold changes between ddPCR and reference datasets in the aforementioned DEGs (Fig. 3c and **Supplementary Table 9**). Note that the level of the correlation coefficients depends on the strength of the intrinsic biological differences. The differences among the Quartet reference materials were relatively small compared to those of the MAQC samples A and B, resulting in relatively lower concordance between the reference datasets and the qPCR or ddPCR data for the Quartet reference materials.

Moreover, we used a LC-MS/MS based proteomics dataset for cross-omics validation of the RNA reference datasets. When all detected genes and proteins were considered, the correlation between RNAseq and proteomics was modest (*r*=0.45∼0.57) (Extended Data Fig. 8), which was similar to what were reported in prior studies (*r*=0.36∼0.60)^34, 35^. However, we found that for differentially expressed genes there was a much higher concordance between RNA and protein data. When using DEGs that were detected by both RNA and protein measurements, the correlation increased to 0.94-0.96 (Fig. 3d and Extended Data Fig. 8). Thus, the protein-coding genes in the reference datasets which were differently expressed in the three sample pairs were successfully validated by the corresponding differential protein abundances. In addition, our findings indicated that the RNA reference datasets might also help benchmark proteomics technologies.

### Reference-dependent quality metrics enhance consistency evaluation between test results and ground truth

To benchmark RNAseq data based on the aforementioned reference datasets, we developed three reference-dependent quality metrics. Specifically, we introduced the “Relative correlation” (RC) metric, i.e., the Pearson correlation coefficient between the fold-changes of a test dataset for a given pair of samples and the corresponding ratio-based reference datasets, representing the trend of numerical consistency of the ratio-based expression profiles. Besides, we introduced the “RMSE” metric, i.e., root mean square error (RMSE) of differences of fold-changes between a test dataset for a given pair of samples and the corresponding ratio-based reference datasets, representing the magnitude of average distances of ratio-based expression profiles. In addition, we introduced the “MCC of DEGs” (MCC) metric, i.e., Matthews Correlation Coefficient (MCC) to measure the consistency of DEGs detected from a test dataset for a given pair of samples with those from the high-confidence DEGs in the reference datasets. Based on their definitions, higher values of RC and MCC of DEGs indicate a better fit between the test dataset and the reference dataset, whereas lower values of RMSE indicate a better fit. All three metrics were able to clearly demonstrate differences in data quality among the 21 batches of data including 13 high-and 8 low-quality batches of data (Extended Data Fig. 9a).

One might argue that the lower RC, higher RMSE, or lower MCC values of the five pre-defined low-quality baches might have resulted from their exclusion during the construction of the reference datasets. To determine whether it was the case or not, we performed a 30-times cross-validation test. Briefly, in each round, we randomly selected 13 batches from the 21 batches to “train” the reference datasets. Reference-dependent quality metrics were then calculated, and the remaining eight batches were used as a “validation” set. The results showed that values of both the “train” and “validation” metrics of the five low-quality batches were consistently lower in RC and MCC or higher in RMSE than those of the 13 high-quality batches, no matter whether they were included or excluded from creating the reference datasets (Extended Data Fig. 9b), demonstrating that the two types of metrics were not dependent on whether the five low-quality batches of data were included in the construction of the reference datasets or not. Instead, the three metrics objectively reflected the intrinsic quality of a dataset, indicating that they were suitable for performance evaluation of future datasets. The cutoff values of RC, RMSE, and MCC values were set to 0.89, 0.38 and 0.54, respectively, which were expressed as the (mean – SD) of RC and MCC, and the (mean + SD) of RMSE across validation sets in the 30-time cross-validation analysis (Extended Data Fig. 9b and **Supplementary Table 2**).

Furthermore, we compared performances between the two categories of quality metrics, including reference-independent quality metric (SNR) and reference-dependent quality metrics (RC and MCC). In most cases, high quality batches showed higher values of SNR, RC and MCC, vice versa, except for one batch (L5_B3) (Extended Data Fig. 9a). In this batch, a high SNR value (16.1) with low reference-dependent quality metrics (RC=0.784, RMSE=0.735, and MCC=0.480) was observed. We found that a customized library preparation kit designed for removing several highly expressed RNAs (e.g., RN7S genes and globin genes) was used in this batch (L5_B3), leading to overall differences between expression profiles from this batch and the reference datasets. Moreover, the complementarity between reference-independent and reference-dependent quality metrics was observed, indicating that both categories of quality metrics should be included in comprehensive performance assessment.

Finally, we calculated a total score by summarizing the two categories of quality metrics. Considering the high correlation among the three reference-dependent metrics (RC, MCC, and RMSE) (absolute *r* > 0.92) (Extended Data Fig. 9a), we used RC to represent reference-dependent metric score for calculating the total score. The total score was expressed as the geometrical mean of SNR and RC for measuring the overall quality of a dataset for the intra-batch proficiency.

### Ratio-based expressions improve cross-batch reproducibility

In large-scale projects, expression profiles are usually measured across multiple batches and pooled together for downstream analysis. Cross-batch reproducibility is therefore crucial. Multi-batch RNAseq datasets derived from the Quartet RNA reference materials allowed us for objective performance assessment of cross-batch reproducibility at multiple levels including cross-time, cross-lab, cross-platform, and cross-protocol.

In this study, after pooling 14 batches of data from the RiboZero protocol together without batch corrections, the impact of batch effects on obscuring the differentiation of biologically distinct groups could be clearly seen in a PCA plot with a diminished SNR value of close to zero (Fig. 4a). Non-experimental factors, rather than intrinsic biological groups (D5, D6, F7, and M8), exhibited the largest differences. When ratio-based expressions were used, which referred to converting expression profiles to gene-wise relative-scale within each batch, the SNR value increased from 0.6 to 22.3. Meanwhile, all libraries from the RiboZero protocol of the same group were grouped together based on ratio-based expressions (Fig. 4b). Similar results were obtained from pooling seven batches of data from the poly(A) protocol and pooling all 21 batches of data from both protocols (Figs. 4a and 4b). These findings indicate the importance of detecting and correcting batch effects in multi-batch studies. Importantly, ratio-based expressions were effective in removing such batch effects.

**Figure 4.**
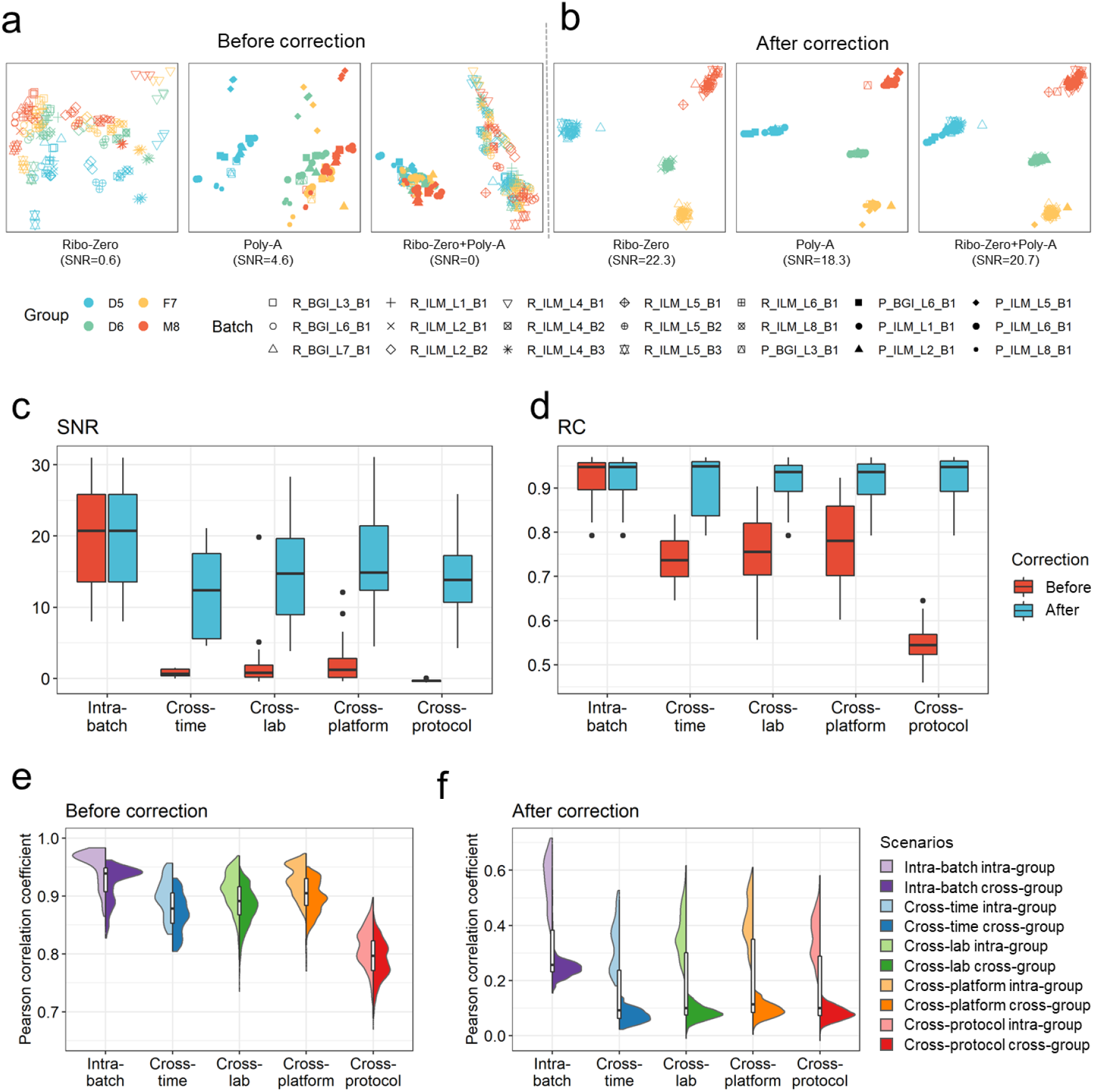
Performance evaluation of cross-batch reproducibility. (**a**-**b**) Scatterplots of PCA on RNAseq data before batch correction (**a**) and after correction (**b**) from replicates of the Quartet RNA reference materials (marked in colors) in the 21 batches (marked in shapes). Expressions in log2FPKM were used as before batch-correction datasets. Ratio-based expressions (which referred to converting expression profiles to gene-wise relative-scale profiles within each batch) were used to correct batch effects. Ratio-based expressions were obtained by subtracting log_2_FPKM by the mean of log_2_FPKM of the three replicates of D6 in the same batch. We used a multi-batch RNAseq dataset, including 168 RNAseq libraries from Ribo-Zero protocol and 84 RNAseq libraries from poly(A) protocol. Plots were color-coded by sample groups and shaped by batches. (**c**-**d**) Boxplots of signal-to-noise ratio (SNR) values (**c**) and relative correlation with reference datasets (RC) values (**d**) for comparisons indicated at the X-axis. When each batch of libraries were compared against each other, they could be classified into five different scenarios with increasing degree of differences, including intra-batch, cross-time, cross-lab, cross-platform of sequencing, and cross-protocol levels. Intra-batch SNR values were calculated using 12 samples in the same batch, whereas SNR values of cross-batch were calculated by combining expression data from all combinations of two batches. (**e**-**f**) Violin-plots of Pearson correlation coefficients based on expression profiles before (**e**) and after (**f**) batch correction for comparisons indicated at the X-axis. D5, F7 and M8 samples were used to calculate pair-wise correlations, while D6 samples were used as denominators for calculating ratio-based expressions for correcting batch effects.

We then compared pair-wise cross-batch performances to investigate integrability at different levels. When each batch of libraries are compared against each other, they could be classified into five different scenarios with increasing degree of differences, including intra-batch, cross-time, cross-lab, cross-platform of sequencing, and cross-protocol levels. We compared the consistency between datasets from different levels of comparison using three quality metrics: SNR, RC, and Pearson correlation coefficients.

SNR values were calculated for the five scenarios of comparisons. Compared to intra-batch SNR values (median SNR = 20.7), SNR values dropped to −0.4∼1.2 (median SNR) at cross-time, cross-lab, cross-platform, or cross-protocol comparisons when absolute expressions (log_2_FPKM) of the two datasets were merged to calculate the SNR value. In this case, it is essentially impossible to distinguish different sample types under the influence of “batch effects”. Thus, expression profiles from two batches of libraries could not be integrated directly at the absolute expression level. However, when ratio-based expressions were used, SNR values maintained as high as 12.4∼14.8 (median SNR) (Fig. 4c). This finding again reinforced the previous notion that ratio-based expressions are more resistant to batch effects (Fig. 4b).

Similar results were obtained for performances based on RC values and Pearson correlation coefficients. Compared to intra-batch RC (median RC: 0.947), RC values dropped significantly to 0.545∼0.780 (median RC) when absolute expressions of two datasets were compared. However, they maintained at 0.936∼0.949 (median RC) when ratio-based expressions of the two datasets were considered (Fig. 4d).

Additionally, the median correlation of absolute expressions was as high as 0.965 for intra-batch technical replicates and 0.937 between different groups in the same batch. It dropped to 0.812∼0.926 for cross-batch technical replicates. What is worse, correlations of technical replicates for the same sample from difference batches are significantly lower than correlations between different sample groups from the same batch (*p* < 0.001), highlighting critical impact of batch effects (Fig. 4e). On the contrary, correlations of ratio-based expressions of technical replicates (0.322∼0.403) were consistently higher than those of different groups (0.072∼0.093) under the different levels of cross-batch comparisons (Fig. 4f), demonstrating the differentiating power at the ratio-based expression level.

Our findings supported the important roles of reference materials in assessing cross-batch reproducibility and their effectiveness in removing batch effects. It should be noted that, we could clearly observe batch effects based on multi-batch datasets of Quartet RNA reference materials (Figs. 4a and 4b), whereas it is impossible with the MAQC reference materials due to their drastic differences (Figs. 1b). Thus, the Quartet reference materials can provide more precise assessment of measurement performance based on their small but biologically relevant intrinsic differences, highlighting their critical roles in assessing cross-batch reproducibility.

### Biological differences between the immortalized B-lymphoblastoid cell lines of the Quartet monozygotic twins

It was noticed that the two immortalized B-lymphoblastoid cell lines corresponding to the two monozygotic twin daughters (D5 and D6) exhibited consistently large differences in gene expression in all batches of data (Extended Data Figs. 5 and 10a), although one might have expected that the expression profiles from the two identical twins would show the highest similarity among all six pairs of the Quartet sample groups. We used ratio-based expression profiles of 13 high-quality batches and applied a weighted gene co-expression network analysis (WGCNA) approach^36^ to discern the underlying biological forces behind the differences in transcriptome between the two cell lines. Genes were grouped with strong co-expression patterns across the sample set into eight modules (Fig. 5a). D5 samples were distinct from D6 samples in the PC1 space based on transcriptomic expression for most modules (7/8), including the largest module (turquoise module) with 2,368 highly co-expressed genes (Figs. 5a and 5b). Functional analysis showed that turquoise-module genes were enriched in Gene Ontology (GO) terms such as cell cycle and B cell proliferation (Fig. 5b). Moreover, a 1,777 gene module (blue module), which showed dispersity between D5 and other three groups (D6, F7 and M8) in the PC1 space, was enriched in B cell mediated immunity. These results implied that differential processes of B cell subtype selection and effects of cell culture might have occurred among Quartet RNA reference materials (Fig. 5b).

**Figure 5.**
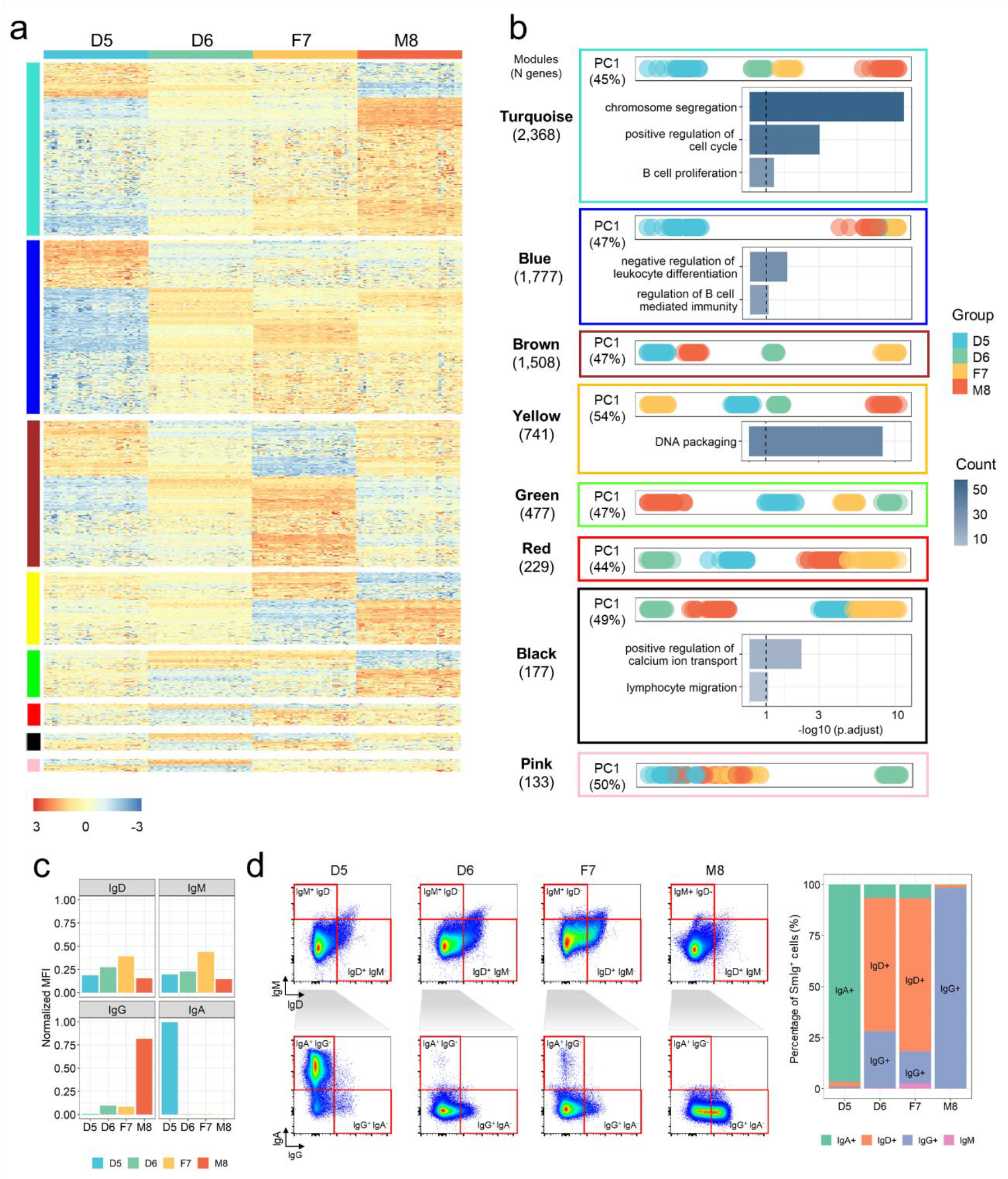
Biological differences between immortalized B-lymphoblastoid cell lines of the Quartet monozygotic twins. (**a**) Expression profiles from four co-expression modules using data from 13 batches with high quality. Color-coded module membership was displayed in the color bars to the left of the dendrograms. Ratio-based expressions were obtained by subtracting log_2_FPKM by the mean of log_2_FPKM of the three replicates of D6 in the same batch. The heatmap was colored using Z-scored ratio-based expression profiles. (**b**) Distances of samples in PC1 space and list of GO terms enriched with genes in each corresponding module. Enriched GO terms were generated using clusterProfiler^55^, with a BH correction and an adjusted *p*-value cutoff of 0.1. PC plots were colored by sample groups. Bar plots were colored based on number of genes included in GO terms. (**c**) The normalized expression level (median fluorescent intensity, MFI) of B-cell surface membrane immunoglobulins (SmIg) IgD, IgM, IgG, and IgA in immortalized LCLs. (**d**) Representative flow cytometric dot plots (left) showed the IgD^+^ cells, IgM^+^ cells, IgG^+^ cells, and IgA^+^ cells in immortalized lymphoblastoid cell lines. (Right) Expression levels of IgA, IgG, IgM or IgD in the four immortalized cell lines.

To identify B cell subtypes corresponding to the Quartet RNA reference materials, we examined the expression levels of B cell surface membrane immunoglobulins (SmIg) on the Quartet cell lines. Four types of Smlgs were measured, including IgD, IgM, IgG, and IgA, which were biomarkers of the developmental stages of B cells. Notably, the IgA expression pattern of the immortalized cell lines from the two monozygotic twin daughters (D5 and D6) exhibited dramatic differences in that IgA was highly expressed in D5 but almost undetectable in D6 (Fig. 5c). Additionally, the expression level of IgG was much higher in the M8 group compared with the other three groups (Fig. 5c). We further performed immunophenotypic analysis of the four immortalized cell lines. In agreement with the SmIg findings from RNAseq, the IgA+ cells were mainly present in the cell line from D5, while a lower percentage of IgA+ cells was found in other cell lines (Fig. 5d). Furthermore, the percentage of IgG+ cells was higher in M8 compared with other three groups (Fig. 5d).

We hypothesized that the major factors driving transcriptomic expression characteristics were probably related to the processes for immortalizing cell lines, e.g., B cell subtype selection during EBV infection and cell culture^37^. To validate this hypothesis, we further conducted RNAseq experiments based on whole-blood samples of the four donors. The twin daughters grouped side-by-side and showed the highest similarity in expression profiles among the Quartet samples (Extended Data Fig. 10b). The intrinsic biological differences between the Quartet monozygotic twins enhanced our understanding of the Quartet RNA reference materials and could be used as another layer of built-in truth to increase the QC utilities of the Quartet RNA reference materials^38^.

### Recommended group-replicate combinations for using the Quartet RNA reference materials for quality control

An important question is what sample-replicate combinations would constitute an appropriate choice for applying Quartet RNA reference materials for quality control in routine transcriptomic profiling. Thus, the replicate number and group number of Quartet reference materials that could be used were enumerated. The results revealed that a minimum of three sample groups and two replicates per batch were required for reaching SNR with high sensitivity for distinguishing data quality of different batches. The use of only two sample groups was not enough for distinguishing quality difference of datasets (Fig. 6a). Meanwhile, it was revealed that a minimum of two replicates per sample type were required for obtaining RC with high consistency with the ground truth (Fig. 6b). Under the same number of replicates and groups, the impact of group combinations (D5, D6, F7, or M8) was minor.

**Figure 6.**
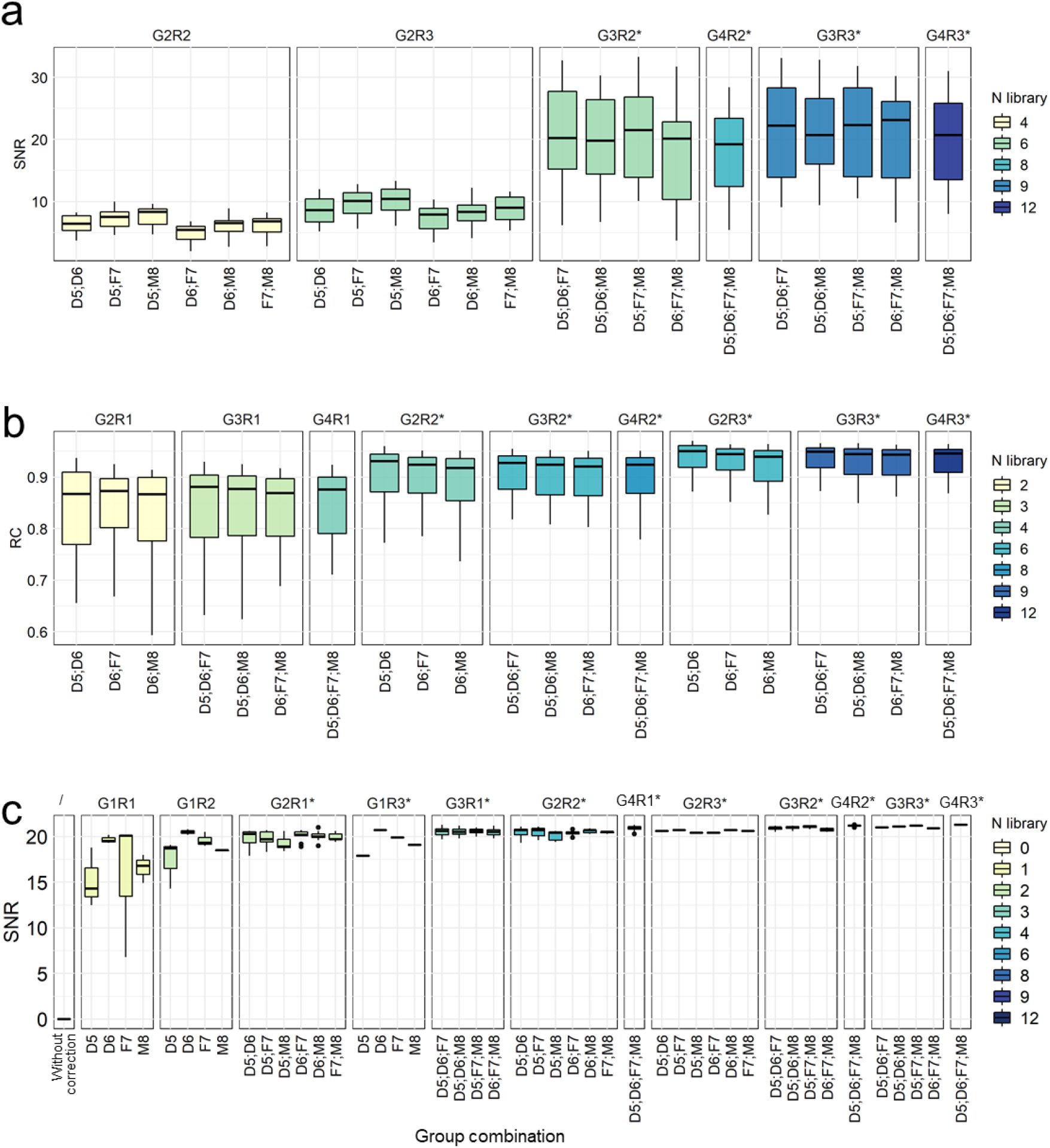
Recommended group-replicate combinations for using the Quartet RNA reference materials for quality control. (**a**-**b**) Distribution of signal-to-noise ratio (SNR) (**a**) and relative correlations with reference datasets (RC) (**b**) under different group-replicate combinations of Quartet RNA reference materials used for assessing intra-batch proficiency. X-axis represented the enumerated number and groups of Quartet reference materials. Titles of sub-panel represented number of libraries, samples and replicates used for SNR calculation. For example, “G2R2” represents four libraries composing two groups (G) with two replicates (R) per group. The recommended combinations were marked with stars (*). (**c**) Distribution of SNR values for ratio-based expression using different numbers of samples and/or replicates as the denominator for the calculation of the ratio-based expressions.

Furthermore, the number and groups of reference materials that could be used as a reference (denominator) in ratio-based profiling within each batch was enumerated. SNR increased dramatically at a ratio (relative) level compared to the absolute level even when only a single replicate was used, with a median SNR value greater than 14 (Fig. 6c). And the SNR values further increased when more replicates and/or more sample groups were added to calculate average expression values as the denominator. Moreover, the SNR values obtained using only one sample to calculate the denominator varied greatly, whereas SNR values based on the mean of more replicates and/or samples as the denominator were stable. Given the same number of samples/replicates as the denominator, a higher number of sample types helped further increase SNR.

These results provided a solid foundation to determine the optimal number of samples and/or replicates to be used for performance assessment and ratio-based transcriptomic profiling using the Quartet RNA reference materials.

## Discussion

We generated large amounts of well-characterized, high quality, homogenous, and stable Quartet RNA reference materials and constructed corresponding reference datasets from reliable transcriptomic data, which can be served as a useful tool for objectively assessing data quality and improve the reliability of transcriptomic profiling.

The Quartet RNA reference materials exhibit several advantages. First, they are established as a part of the multiomic reference materials, with matched DNA, RNA, protein, and metabolite generated from the same immortalized cell lines. This study design allows for cross-omics validation and will help reliably understand the biological traits of the reference materials. Secondly, the suite of reference materials are from a four-member Quartet family including two monozygotic twin daughters and their father and mother. Genomic and phenotypic characteristics are involved in the four RNA samples, acting as built-in “truths”. Reference datasets based on intrinsic biological differences among the Quartet RNA reference materials have been constructed and can be used as “ground truth” for quality assessment (Fig. 3). Expression characteristics affected by genetic relationships of the four reference materials will be further studied^38^. Thirdly, the RNA reference materials are derived from cell lines from four individuals. The small intrinsic biological differences among the Quartet reference materials enable precise assessment at inter-patient level, rather than inter-tissue level, which are closer to clinical scenarios (Fig. 1c). Fourth, the RNA samples have been produced in large amounts in one batch and are renewable through cell culture. By minimizing batch effects that may be introduced during cell culture and RNA extraction, the Quartet RNA reference materials are sufficient for performing standard RNAseq experiments over 10,000 to 50,000 times and provide a material basis for long-term quality monitoring. Based on comprehensive assessments, the Quartet RNA reference materials are homogenous and long-term stable at the storage temperature of −80°C. The publicly available Quartet RNA reference materials can also be used for further evaluation of emerging technologies, as well as new areas of interest which are beyond gene expression levels, such as alternative splicing, RNA-editing, gene fusion, and epi-transcriptomics. In addition, the Quartet RNA reference materials comprise high-quality total RNAs, including not only full-length RNAs, but also small RNA molecules such as miRNAs, enabling further quality assessment of small RNA profiling technologies.

Quality metrics derived from the Quartet RNA reference materials and reference datasets can be used for proficiency testing and external quality assessment. Previous quality metrics were focused on biases from library preparation, or detecting outliers in expression profiles^1, 15, 19, 39–42^. The MAQC/SEQC consortia have demonstrated repeatedly that “lab effects” strongly affect the detection of DEGs, highlighting the importance of assessing data quality in detecting DEGs^18^. In this study, we developed comprehensive quality metrics for assessing the reliability of differential expression, including discriminating power across different biological groups (SNR) and reproducibility of identifying DEGs (RC and MCC), which are reference-independent and reference-dependent, respectively. In addition, distributions of these QC measures were obtained from multiple real-world RNAseq datasets, providing practical cutoffs to decide whether the proficiency of a test dataset is acceptable.

The Quartet RNA reference materials can be used for monitoring and correcting batch effects. Batch effects are notorious technical variations irrelevant to study factors and are challenging to deal with, especially when they are confounded with biological factors of interest^43, 44^. Our results demonstrated that the presence of batch effects without correction can lead to misclassification of samples (Fig. 4a). Fortunately, the batch effects can be mitigated by using ratio-based expressions (Figs. 4b-4f) if one or more common reference materials are profiled across batches. Our companion work has found that “ratio” data by scaling the absolute feature values of study samples relative to those of concurrently measured reference sample(s) on a feature-by-feature basis could effectively mitigate the widespread problems of batch effects, in transcriptomics, proteomics, and metabolomics datasets^45^. Importantly, the “ratio” method is equally effective even for study design of unbalanced distributions of samples in different groups between different batches. In practice, the imbalance in samples across batches is almost inevitable because of hidden biological subpopulation variabilities^43, 44^. Therefore, the “ratio” profiling based on Quartet RNA reference materials in each batch can play an important role in making expressions more reproducible and resistant to batch effects.

The Quartet RNA reference materials can act as valuable tools for quality control in large-scale, longitudinal, and multi-center projects. Many large-scale consortium projects with comprehensive and coordinated efforts help accelerate our understanding of the molecular basis of transcriptome by producing RNAseq data with a large sample size^31, 46–48^. However, the broad variety of platforms, protocols, and lab proficiencies^49–51^ has created the need for comprehensive reference materials. At the starting point of a large-scale project, we recommend researchers conducting RNAseq experiments using the Quartet RNA reference materials in each lab to assess and ensure intra-batch proficiency and cross-batch reproducibility before analyzing precious study samples. Meanwhile, researchers can use the Quartet RNA reference samples routinely along with study samples to monitor and correct batch effects.

The combinations of sample types and number of replicates for the application of the Quartet reference RNAs are context dependent. For proficiency test and external quality assessment purposes, where the frequency of reference usage could be as low as a few times per year, it is recommended to apply multiple types of samples with multiple replicates per sample. Users can apply a minimum of three sample groups and two replicates for quality assessment (Fig. 6). Without too much concern on cost, users can apply a total of 12 samples, comprising the four Quartet RNA samples with three replicates for each RNA sample, to implement full quality assessment mentioned in the study and remove batch effect in a robust way. For batch effect removal purpose in large cohort studies, where the Quartet reference RNA samples are expected to be routinely used along with study samples and additional cost associated with profiling reference samples becomes an issue, it is recommended to apply fewer sample types and fewer replicates per batch. Users can even apply four sample types or as few as two sample types without replicates as a cost-effective choice of references for monitoring and correcting batch effects. In this case, we suggest the use of a total of four profiles from each replicate of the four Quartet RNA reference materials as the denominator per batch of 96 libraries for ratio-based expression profiling, reaching a high SNR while maintaining a reasonable additional cost (4/(96-4)=4.3%) per batch of 92 study samples.

To facilitate the adoption of multiomic reference materials, reference datasets, and quality metrics from the Quartet project, we developed a Quartet Data Portal (http://chinese-quartet.org/) for access to the Quartet resources and enhancing quality consciousness of the community, as described in an accompanying paper^52^. Researchers can request the multiomic reference materials, multi-level datasets, and reference datasets from the data portal. Besides, researchers can upload Quartet RNAseq data of their own, automatically run analysis to evaluate data quality, and/or share data with the community. With the growing use of the Quartet reference materials, we hope to generate and collect diverse datasets to facilitate upgrades of the reference datasets and quality metrics.

Though many advantages of using the Quartet RNA reference materials were obvious and listed above, several limitations of the Quartet samples should also be noted. First, only around 52∼54% of the 58,395 annotated genes were reliably detected in the Quartet RNA reference materials, limiting quality assessment and ratio-based scaling to these detectable genes. Secondly, a single analysis pipeline was used in this study, which may introduce bias in transcriptomic quantification and characterization of the reference datasets. Although prior studies have compared the performance of different RNAseq analysis tools and found overall good reproducibility for different tool combinations in terms of differential expression calls after proper filtering processes^53, 54^, bioinformatics tools will be further evaluated and used for characterizing the reference datasets. Thirdly, the datasets were generated by high-throughput short-read sequencing technologies. With widespread adoption of reference materials and routine reference benchmarking, we encourage the “bake-offs” among labs and platforms for evaluating additional reagents, protocols, and instruments.

In summary, the Quartet RNA reference materials and reference datasets are unique resources to improve quality of RNAseq data. Inclusion of the Quartet RNA reference materials in RNAseq batches coupling with reference datasets will make RNAseq more reproducible, accurate, and comparable, and therefore improve clinical applications and biomedical research of transcriptomics.

## Materials and Methods

### Cell lines

Human subjects, establishment of the Epstein-Barr virus (EBV) transformed B-lymphoblastoid cell lines, expansion and cryopreservation of the cells, cell culture and cell quality control (QC) have been described in an accompanying paper by Zheng et al^29^. Briefly, four healthy volunteers from a quartet family in Taizhou, Jiangsu, China were enrolled, and their peripheral blood samples were collected. The study was approved by the IRB (Institutional Review Board) of the School of Life Sciences, Fudan University (BE2050). Peripheral blood mononuclear cells (PBMC) were isolated, and the naïve B cells were sorted and infected with EBV by centrifugation at 2000 rpm for one hour and the immortalized cell lines were cultured in an incubator. About 1.0×10^11^ cells were harvested for each cell line in the same batch to ensure that multiomic reference materials were extracted from the same batch of cultured cells. About 2.0×10^9^ cells per cell line were used for generating Quartet RNA reference materials.

### RNA extraction and quality assessment

RNA reference materials were extracted from EBV immortalized B-lymphoblastoid cell lines. TRIzol reagent was added to resuspend the cells. Total RNA was extracted using RNeasy Maxi kit (Qiagen GmbH, Germany, cat#: 75162) including on-column DNase-I digest, according to the manufacturer’s instructions. Total RNAs including miRNAs and other small RNA molecules were retained, allowing for both RNA and small RNA profiling.

The RNA integrity number (RIN) values were obtained for assessing RNA quality with a Bioanalyzer 2100 (Agilent Technologies) using RNA 6000 Nano assay (Agilent Technologies) and a Qsep 100 system (BiOptic Inc., Taiwan, China). RNA concentrations, OD280/260, OD260/230, 28/18S were assessed by NanoDrop ND-2000 spectrophotometer (Thermo Scientific). Over five milligrams of RNA were obtained per cell line. RNAs were then aliquoted into more than 1000 tubes per sample group with 5 μg of RNA per tube.

As a part of the Quartet Project, multiomic reference materials (DNA, RNA, protein, and metabolite) were established simultaneously from the same batch of cultured EBV immortalized B-lymphoblastoid cell lines from the Quartet family members. The establishment of Quartet multiomic reference materials have been described in an accompanying paper by Zheng et al^29^. The Quartet multiomic reference materials are available to the public. Users can request all types of reference materials via the Quartet Data Portal (http://chinese-quartet.org/).

### RNA stability assessment

#### Bottle-opening and freeze-thaw stability

RNAs were stored in 0.5 mL tubes at −80 °C for over 1 h until completely frozen. Frozen samples were thawed at 4 °C for approximately 0.5 hrs until completely thawed (freeze-thaw 1). We then opened the tubes and took 1 μL aliquots per tube out for further analysis (bottle-opening 1). The remaining RNAs were immediately re-frozen at −80 °C. This cycle was repeated for 20 times. RIN values were assessed at the 0, 1, 2, 3, 4, 5, 6, 8, 10, 14, 16, 18 and 20 times of opening and freeze-thaw to evaluate the integrity of RNA. Three replicates per sample group were assessed during each assessment.

#### Short-term stability

The stability of Quartet RNA reference materials at room temperature (22 ∼ 25 °C) and 4 °C was assessed. First, four groups of the Quartet RNA reference materials were assessed for up to four days. RIN values were assessed at 0 hr, 6 hrs, 24 hrs and four days to evaluate the overall quality of RNA during storage. Secondly, considering the same trends and similar results across the four Quartet RNA reference materials, we used two RNA reference materials (F7 and M8) for up to 14 days. RIN values were assessed at 0, 2, 4, 5, 6, 7, 8, 10,12,14 days separately. Three replicates per sample group were assessed during each assessment.

#### Long-term stability

The stability of RNA reference materials at storage of −80 °C was monitored for up to 20 months. RIN values were assessed at the 0, 1, 2, 3, 4, 5, 6, 7, 8, 9, 11, 12, 13, 15, 16, 17, and 20 months. Three replicates per sample group were assessed each timepoint. The MAQC/SEQC RNA reference materials, including A sample (Universal Human Reference RNA, Agilent Technologies, Inc.) and B sample (Human Brain Reference RNA, ThermoFisher, Inc.)^13^, were used as controls at each timepoint.

### Library construction and sequencing

According to the Quartet Project study design, 12 tubes of RNA samples were sent to each lab, including four groups of the Quartet RNA reference materials with triplicates per group. Library preparation, library QC, and sequencing were conducted in a fixed order (D5-1, D6-1, F7-1, M8-1, D5-2, D6-2, F7-2, M8-2, D5-3, D6-3, F7-3, and M8-3) in each lab, to eliminate confounding factors such experimental sample-processing order with sample group.

RNAseq library preparation and high-throughput sequencing were conducted by each lab. Briefly, libraries were constructed by poly(A) selection or ribosomal RNA depletion (RiboZero) methods. The libraries were sequenced on Illumina NovaSeq or BGI BGI2000 platforms with paired-end (PE) reads of 100-150 bp. A total of 252 Quartet RNAseq libraries from 21 batches were generated. Additionally, we simultaneously generated 20 batches of RNAseq datasets using MAQC reference materials (MAQC A and B samples) as controls, enabling head-to-head comparisons of the two sources of RNA reference materials. Detailed information on RNAseq library construction and sequencing was shown in **Supplementary Table 1**.

Four RNA libraries from whole blood of the Quartet donors were constructed by ribosomal RNA depletion methods (TruSeq RNA Library Prep Kit) and sequenced on Illumina HiSeq 4000 platform with 150 bp PE reads.

### Alignment and gene quantification

Preliminary processing of raw fastq reads was performed using fastp v0.19.6 to remove adapter sequences^56^. Read alignment and quantification was conducted using HISAT v2.1, SAMtools v1.3.1, StringTie v1.3.4 and Ballgown v2.14.1^57^. Reference human genome build 38 (https://genome-idx.s3.amazonaws.com/hisat/grch38_snptran.tar.gz) and gene model from Ensembl (http://ftp.ensembl.org/pub/release-93/gtf/homo_sapiens/Homo_sapiens.GRCh38.93.gtf.gz) were used for read mapping and gene quantification. log2 transformation was then conducted based on Fragments Per Kilobase of transcript per Million mapped reads (FPKM) values. To avoid infinite values, a value of 0.01 was added to the FPKM value of each gene before log2 transformation. Expression profiles based on detected genes were used for further analysis. A gene was considered detectable (expressed) in a biological group within a batch if ≥ 3 reads were mapped onto it in at least two of the three replicates.

Quality control analysis of sequencing data at pre-alignment and post-alignment level was conducted using FastQC v0.11.5^58^, FastQ Screen v0.12.0^59^, Qualimap v2.0.0^60^, and MultiQC v1.8^61^.

### Differentially Expressed Genes (DEGs)

Differential expression analyses were implemented using the limma v3.38.3^62^ and edgeR v3.24.3^63^ packages according to guidelines from the limma package. Briefly, after filtering low-expressed genes, read counts of the kept genes were normalized with the trimmed mean of the M-values normalization (TMM) method. The voom function was then employed to estimate the mean-variance relationship. Functions of lmFit, contrasts.fit, and eBayes were applied to estimate the fold change of each gene between the two sample groups with a p-value. A gene was considered differentially expressed in a batch between two sample groups if p < 0.05 and fold change ≥ 2 or ≤ 0.5 using limma package for up- or down-regulation, respectively.

### Identification and quantification of alternative splicing

The alignment results based on the HISAT2 were used to conduct the alternative splicing (AS) graphs, identify and quantify AS events, using SplAdder toolkit^64^. Briefly, we used SplAdder with the parameters of confidence level 2 for generating individual graph of each sample and merge them into a single comprehensive AS graph for all samples later. Based on the graph, SplAdder was used to quantify six types of AS events, including exon skip, intron retention, alternative 3’ splice site, alternative 5’ splice site, cassette exon, and coordinated mutually exclusive exons. The percent spliced in (PSI) values based on the quantified splicing graphs for each event was used for further analysis^65^.

### Construction of reference datasets

We constructed the reference datasets of ratio-based expression based on the following steps: (1) identifying detectable genes; (2) calculating ratio-based expression based on reliable genes which were differentially expressed and with low uncertainty; (3) assessing the homogeneity and stability; (4) assessing the uncertainty of ratio-based reference datasets; and (5) calculating high-confidence differentially expressed genes (DEGs) in reference datasets.

First, detectable genes were identified. A gene was considered expressed in a sample in each batch if more than three reads were mapped to it in at least two of the three replicates. If a gene was detected in all the 13 batches in a sample group (D5, D6, F7 and M8), it was considered expressed in that sample group.

Secondly, ratio-based expressions were calculated. We used the expression profiles of three replicates of D6 in the same batch as the denominators and derived the ratio-based expressions for the three sample pairs (D5/D6, F7/D6, and M8/D6). The reference ratio-based expressions between each pair of samples for a gene was provided in the format of an average by summarizing from the 13 fold-changes calculated from each of the 13 high-quality RNAseq datasets. In order to improve the reliability of the reference values, genes were included if they satisfied the following criteria: (1) Detectable across the two groups of each sample pair; (2) Limma-based^62^ *p* < 0.05 in at least four batches in each sample pair; (3) not significantly different between poly(A) and RiboZero protocols (student t-test *p* > 0.05 or fold change < 2 and ≤ 0.5).

Thirdly, the homogeneity and stability was assessed using RNAseq datasets. The Quartet RNA reference materials were considered to be homogenous and stable, as can be seen from the corresponding reference datasets. Additionally, uncertainties of reference materials were assessed.

Finally, high-confidence DEGs in the reference datasets (reference DEGs) were identified. A gene was considered as a reference DEG between two sample types if it was concordantly discovered as an up- or down-regulated gene (*p* < 0.05 and fold change ≥ 2 or ≤ 0.5) in more than six of the 13 high-quality batches.

### Homogeneity assessment based on RNAseq datasets

The homogeneity of the Quartet RNA reference materials was assessed using RNAseq data. We randomly selected 17 tubes (units) of each Quartet RNA reference material and named them as N1-N17. Under the same condition, nine replicates in N1 tube and one replicate in tube N2-N17 of each material were assessed to represent within-unit (n=9) and between-unit (n=16) characteristics (Extended Data Fig. 7a). A total of 25 RNAseq experiments per reference material were conducted.

RNAseq libraries were constructed by ribosomal RNA depletion methods (VAHTS^®^ Universal V6 RNAseq Library Prep Kit for Illumina) and sequenced on the Illumina NovaSeq platform with 150 bp PE reads. Alignment, quantification, and quality control were conducted using the same analysis pipeline and parameters described above.

The within-unit and between-unit variances were then calculated using the ANOVA method^32, 33^. Ratio-based expressions were obtained by subtracting log_2_FPKM by the mean of log_2_FPKM of the three replicates of D6 in the same batch and used. A gene was considered to be homogeneous when a cutoff of FDR-adjust ANOVA-based *p* >0.05 was used. Only between-unit homogeneity is studied, since within-unit homogeneity might be negligible in the case of intrinsically homogeneous materials such as solutions^66^.

### Long-term stability assessment based on RNAseq datasets

We assessed the long-term stability of the reference materials 15 batches of RNAseq datasets which were generated from up to 26 months (Extended Data Fig. 7c). Ratio-based expressions were obtained by subtracting the mean log_2_FPKM of the three replicates of D6 in the same batch from the log_2_FPKM values. According to ISO Guide 35 (2017)^32^ and JJF-1343 (2012)^33^, long-term stability assessment was conducted based on regression analysis. For each gene, the observed slope *b*_1_ and uncertainty of slope *b*_1_ (*s*(*b*_1_)) was calculated. If |*b*_1_|< s(*s*(*b*_1_)) × t_0.95,n-2_, the expression of the gene is stable, vice versa, where t_0.95,n-2_ is critical *t* value for a confidence level of 95% and *n*-2 degree of freedom.

### Uncertainty assessment of reference datasets

According to ISO Guide 35 (2017)^32^ and JJF-1343 (2012)^33^, the source of uncertainties can be classified into characterization uncertainties (*u_char_*), sample inhomogeneities (between-bottle variation, *u_bb_*), and instabilities (*u_s_*). These values were then combined to combined uncertainties (*u_c_*) and expanded uncertainties (*U*) with an expansion factor.

First, characterization uncertainty of genes in the reference datasets was evaluated, using 13 fold changes (log2 scale) from each of 13 high-quality RNAseq datasets. Relative uncertainty of characterization was used as characterization uncertainty (*u_char_*), which can be expressed as E.q. (1) as follows:

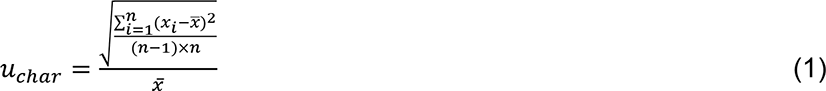

where *n* is number of measurements in the sample; *x*_*i*_ is measurement value of i^th^ time; *x̅* is average value of *x* across n times.

Secondly, sample inhomogeneity *u*_*bb*_ was evaluated using RNAseq datasets s (Extended Data Fig. 7a). *u*_*bb*_can be expressed as equation as E.q (2) and (3):

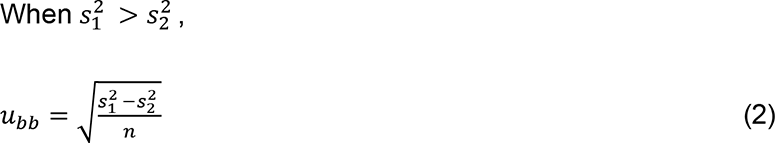

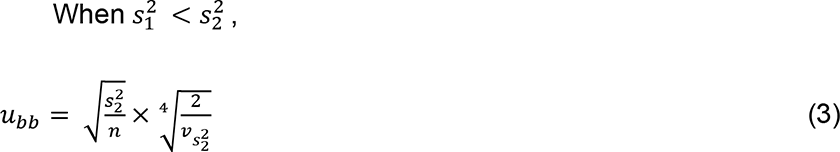

where 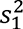 is between-unit variation; 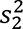 is within-unit variation; 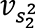 is degree of freedom of 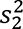; *n* is number of between-unit measurements.

Thirdly, long-term instability (*u*_*s*_) was evaluated based on RNA quality RIN across 20 months, which can be expressed as E.q. (4)

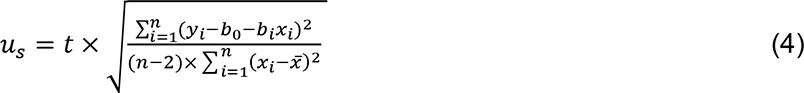

where b_0_ and b_1_ is the intercept and slope of linear regression line between x_i_ (month) and y (RIN); t is time (month); n is number of observations. Short-term instability might be negligible, since reference materials are recommended to be transported using dry ice.

Fourthly, a combined uncertainty (*u*_*c*_) should consider all uncertainty described above, which can be expressed as E.q. (5):

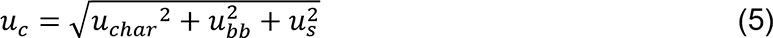

Finally, an extended uncertainty (*U*) can be expressed as E.q. (6):

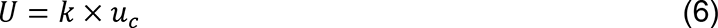

where k is a constant value. Here, *k* = 2 was applied for 95 % confidence level.

### Performance metrics

Performance metrics including signal-to-noise ratio (SNR), relative correlation with reference datasets (RC), root mean square error (RMSE) of differences with reference datasets, and Matthews Correlation Coefficient (MCC) of DEGs were developed to evaluate the quality of RNAseq data at expression level before a total score was calculated.

#### SNR

Signal-to-noise ratio (SNR) is a measurement used in science and engineering. SNR is defined as the ratio of the power of a signal to the power of noise and is often expressed in decibels (https://en.wikipedia.org/wiki/Signal-to-noise_ratio). In this study, the average distances representing the intrinsic “differences” among distinct biological sample groups are regarded as the signal, whereas the average distances among technical replicates of the same sample group are regarded as noise.

To identify an effective way to characterize the SNR values, we evaluated the performances of SNR values calculated by different algorithms, including Euclidean distance (Dist), overall expression profiles (Expr), Pearson correlation coefficient (Cor), t-Distributed Stochastic Neighbor Embedding (tSNE), and principal components analysis (PCA), and found PCA based SNR outperformed. The numbers of PC components used in calculating SNR were then determined. We decided to use the first two components in PCA to calculate SNR values in correspondence with visualization in PCA plots.

Therefore, SNR is defined as E.q (7):

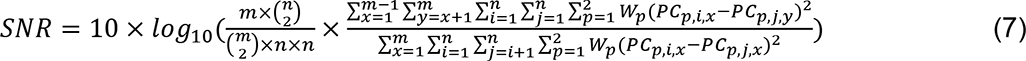

where *m* is the number of sample groups, while *n* is the number of replicates in each sample group. *W*_*p*_ represents the *p*^th^ principal component of variances. *PC*_*p*,*i*,*x*_, *PC*_*p*,*j*,*x*_ and *PC*_*p*,*j*,*y*_ represent the *p*^th^ component values of replicate *i* and replicate *j* in sample group *x* or sample group *y*, respectively.

A standard sample set consisted of 12 tubes with each representing one of the three replicates of the four RNA reference materials. Therefore, a typical SNR in the study was the ratio of the average distances between different biological groups (9*12/2=54) to the average distances between technical replicates of the same groups (2*3*4/2=12). The distribution of intra-batch SNR values from 21 RNAseq datasets was used to identify a threshold of 12 (mean-standard deviation), indicative of high discriminating power.

#### RC

Relative correlation with reference datasets was calculated based on the Pearson correlation coefficient between the ratio-based expression levels of a dataset for a given pair of groups and the corresponding reference fold-change values. It is referred to as the “relative correlation with reference datasets” metric, representing the numerical consistency of the ratio-based expression profiles. To improve reliability, the mean of the three replicates of each sample group was calculated before performing ratio-based expression analysis. Fold-changes were transformed using log2 scaling.

#### RMSE

RMSE was calculated using fold-changes between a test dataset for a given pair of samples and the corresponding ratio-based reference datasets, representing the average distances of ratio-based expression profiles. Fold-changes were transformed using log2 scaling. It was implemented using *rmse* function from *Metrics* package^67^.

#### MCC

MCC is a widely used statistic in the field of bioinformatics and machine learning, which combines test sensitivity and specificity^18, 68^. In this study, we used MCC to measure the consistency of DEGs detected from a dataset for a given pair of samples with those from the reference DEGs, or “MCC of DEGs”. Reference DEGs and non-DEGs as true positive and true negative sets were integrated by consensus voting. When DEGs and non-DEGs of a given dataset were identified, the number of True Positive (TP), True Negative (TN), False Positive (FP) and False Negative (FN) could be calculated. MCC is computed using the E.q. (8):

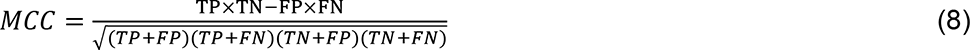

#### Total score

The total performance score is calculated to measure the overall quality of a dataset generated from a lab for its effectiveness in quantifying the transcriptomic differences among the four Quartet RNA reference materials by summarizing reference dataset-independent quality measurement (SNR) and reference dataset-dependent quality measurement (RC). The total score is expressed as the geometrical mean of SNR and RC.

### Cross-validation of reference-based quality metrics

To examine if the lower RC, higher RMSE, or lower MCC and MCC for the five low-quality batches were caused by their exclusion from creating the reference datasets, we performed 30 times of cross-validation test. Briefly, in one cross validation, we randomly selected 13 batches from the 21 batches to create (“train”) the reference datasets which were then used to calculate quality measurements for all the 21 batches. Both high- and low-quality batches might be randomly included or excluded from “training the reference datasets”, either as training or validation sets.

### Co-expression analysis

Co-expression network was constructed using the WGCNA (v1.71) package ^36^ using ratio-based expression profiles of 13 batches with high quality. Ratio-based expressions were used by scaling FPKM by the geometric mean of FPKM of the three replicates of D6 in the same batch. Genes with the highest variations (n=10,000) were used for conducting co-expression network. Average linkage hierarchical clustering was performed to group genes based on topological overlap dissimilarity measurement of their network connection strengths. Modules were then identified with a dynamic tree-cutting algorithm with a minimum module size of 50. Soft power was set to 6. Modules were named in color. Eight modules were identified, including turquoise (n=2,368), blue (n=1,777), brown (n=1,508), yellow (n=741), green (n=477), red (n=229), black (n=177) and pink (n=133). The remaining genes (n=) did not fit elsewhere. PCA and functional analysis of each module were conducted.

### RT-qPCR

Primers of 83 genes were designed using online Primers-BLAST of NCBI based on the RNA sequences. And the PCR method of reference gene (C1ORF43) was established previously. Primers were synthesized by Beijing Liuhe Huada Gene Technology Co., Ltd (**Supplementary Table 8**).

RT-qPCR reactions were performed in two steps. First, reverse transcription was carried out using 2 μL of RNA mixed with 4 μL of 5×PrimeScript IV cDNA Synthesis Mix (Takara, Code No. 6215A) containing PrimeScript IV RTase, RNase Inhibitor, Oligo dT Primer, dNTP Mixture buffer and 1 μL Random 6 mers, and nuclease free water up to 20 μL final reaction volume. This reaction mixture was incubated at 30°C for 10 minutes, then for 15 minutes at 42 °C and finally for 5 minutes at 95°C for termination. Secondly, cDNA obtained in the previous step was used as template for qPCR. The qPCR reactions were carried out using UltraSYBR Mixture (Low ROX) (CWBIO, CW2601M) containing 2 μL of cDNA, 0.4 μL of each forward and reverse primers (final concentration of 200nM) in a 20 μL final volume reaction. The qPCR was performed on a Roche 480 qPCR System using the following cycling conditions: 10 minutes at 95°C followed by 45 cycles of 15 seconds at 95 °C and 1 minute at 60 °C. Three replicates per sample per gene were conducted for eliminating random variations.

Comparative Ct method (delta delta Ct method) was used to calculate the fold differences for the three sample pairs (D5/D6, F7/D6 and M8/D6) with housekeeping gene C1ORF43 as endogenous control. For the qPCR data, a gene is called DEG when the student’s t-test *p*-value < 0.05 and fold change ≥ 2 or ≤ 0.5.

### ddPCR

DEGs identified in reference datasets and qRT-PCR were further validated using droplet digital PCR (ddPCR). The ddPCR reaction was performed in a QX200 Droplet Digital PCR System (Bio-Rad Laboratories, CA) according to the manufacturer’s instructions. Each test was prepared in a total of 20 μL volume of the reaction mixture, comprising 10 μL EvaGreen® Supermix (Bio-Rad), 2 uL of forward and reverse primers, 2 μL of cDNA templates, and 6 μL of RNase Free ddH_2_O. Samples and 70 μL droplet generation oil were then placed into Droplet Generator (Bio-Rad). Droplets (40 μL) were transferred to a 96-well PCR plate. The PCR reactions were performed using the following cycling conditions: Pre-denature for 1 cycle at 95 °C for 5 min; denature for 40 cycles at 95 °C for 30 s; anneal and extend for 40 cycles at 60 °C for 1 min. After the cycles, a signal stabilization step of 4°C for 5 min and 90°C for 5 min was conducted. The signals were read by Droplet Reader (Bio-Rad). Each reaction was performed in duplicate.

### Flow cytometry

Immortalized B-lymphoblastoid cell lines were harvested, washed, and stained with fluorescence-conjugated antibodies. The following antibodies were used for cell surface staining: IgA (clone IS11-8E10) (Miltenyi Biotec, cat#: 130-114-002), IgD (clone IA6-2) (BD Biosciences, cat#: 561314), IgG (clone G18-145) (BD Biosciences, cat#: 561296), and IgM (clone G20-127) (BD Biosciences, cat#: 562977). Flow cytometric analyses were performed on CytoFLEX LX (Beckman Counter), and data were analyzed with FlowJo V10.7.2 software (BD Biosciences).

### LC-MS/MS based proteomics

Mass Spectrometry (MS)-based data-dependent acquisition (DDA) proteomics datasets from Quartet protein reference materials was used for cross-omics validation. Detailed description of sample preparation, data generation was provided by Zheng et al^29^. Briefly, large quantities of Quartet peptide reference materials (Lot: 20200616) were generated from LCLs. LC-MS/MS based proteomics datasets (4 groups × 3 replicates) were then generated in a lab (code: NVG) using Q Exactive HF-X mass spectrometer (ThermoFisher). To eliminate confounding factors such as experimental sample-processing order with sample group, experiments were conducted in a fixed order (D5-1, D6-1, F7-1, M8-1, D5-2, D6-2, F7-2, M8-2, D5-3, D6-3, F7-3, and M8-3).

Peptide and protein identification and quantification was conducted using Proteome Discoverer 2.2 (PD 2.2, ThermoFisher) based on the human reference database UniProt (http://www.uniprot.org). Proteins with at least one unique peptide with 1% FDR at the peptide level were retained for further analysis. Protein quantification was normalized using the fraction of total (FOT). The FOT was multiplied by 10^5^ for the ease of presentation.

### Biological classifications from published datasets

We used publicly available datasets to examine the extent of biological differences with four “intrinsic” biological classification groups from published datasets^7, 30, 31^. Jiang et al.^7^ characterized primary Chinese triple-negative breast cancer (TNBC) patients and classified them into four transcriptome-based subtypes (BLIS, LAR, IM, and MES). Expression profiles of the four molecular subtypes of breast cancer were downloaded from the Genomic Data Commons (GDC) Data Portal^30^, including Luminal A (N=420), Luminal B (n=174), basal-like (n=140), and Her2 enriched (n=65). Expression profiles from four cancer types with distinct tissue types were also downloaded from the GDC Data Portal^30^, including brain cancer (n=80), breast cancer (n=80), kidney cancer (n=80), and lung cancer (n=80). Expression profiles from four normal tissue types were obtained from GTEx, including brain (n=100), liver (n=100), lung (n=100), and muscle (n=100)^31^. A gene was considered as a DEG when t-test *p* < 0.05 and fold change ≥ 2 or ≤ 0.5.

### Statistical analysis

All statistical analyses were performed using R statistical packages (v3.6.1 and v4.1.2) (https://www.r-project.org). PCA was conducted with the univariance scaling, using the prcomp function. Hierarchical clustering analysis (HCA) was performed using Ward linkage based on a distance matrix using Euclidean method to measure the distance of the samples and genes, using R package pheatmap v1.0.12 (https://rdrr.io/cran/pheatmap/). Differential expression analyses were implemented using the limma v3.50.0 and edgeR v3.36.0 packages according to guidelines from the limma package^62, 63^. Functional enrichment analyses based on GO terms were conducted with the R/Bioconductor package clusterProfiler v4.2.2, with a BH correction and an adjusted *p*-value cutoff of 0.05 ^55^. Data visualization was implemented using R package ggplot2 v3.3.5 (https://ggplot2.tidyverse.org/), GGally v2.1.2 (http://ggobi.github.io/ggally/), and ggsci v2.9 (https://github.com/nanxstats/ggsci). The workflow was created with BioRender.com.

### Data availability

According to the regulation of the Human Genetic Resources Administration of China (HGRAC), data produced in this study are available at NODE OEP000970 (https://www.biosino.org/node/project/detail/OEP000970) and the Genome Sequence Archive (GSA) of the National Genomics Data Center of China with BioProject ID of PRJCA007703. Moreover, we have developed the Quartet Data Portal (http://chinese-quartet.org) for the community to access and share the Quartet multiomic resources.

## Disclaimer

The content is solely the responsibility of the authors and does not necessarily represent the official views of the US Food and Drug Administration. This research was supported in part by the Intramural Research program of the National Library of Medicine, US National Institutes of health.

## Acknowledgments

This study was supported in part by National Key R&D Project of China (2018YFE0201603 and 2018YFE0201600), the National Natural Science Foundation of China (31720103909 and 32170657), Shanghai Municipal Science and Technology Major Project (2017SHZDZX01), State Key Laboratory of Genetic Engineering (SKLGE-2117), and the 111 Project (B13016). Some of the illustrations in this paper were created with BioRender.com.

## Author contributions

Y.T.Z., R.Z., F.Q., J.X., Y.Y., L.S., L.J., X.F., J.L., W.T., H.H., and W.X. conceived and oversaw the study. Y.T.Z., W.H., H.W., S.S., Z.C., P.Z., Y.Z., R.L., S.Z., X.W., D.B., and B.Y.L. cultured the cell lines, prepared and characterized the RNA reference materials. J.G. and F.Q. performed flow cytometry assays and data interpretation. Y.T.Z., W.H., L.D., H.W., J.H., and S.S. coordinated and/or performed NGS library preparation and sequencing. L.D., Y.T.Z., X.W., Y.P., and Y.Y. performed qPCR validation. Y.Y., W.H., H.W., L.D., Y.L., S.S., J.Y., Z.C., Q.C., Z.L., Z.C., N.Z., J.L., L.R., H.J., J.S., T.Q., B.L., C.S., F.D., A.S., P.M., J.H., L.Z., H.J., D.T., J.T., W.X., H.H., W.T., J.W., J.L., X.F., L.J., L.S., J.X., F.Q., R. Z., and Y.T.Z performed data analysis and interpretation. Y.Y., J.Y., and J.S. managed the datasets. Y.Y., Y.T.Z., R.Z., F.Q., J.X., W.T., H.H., W.X., and L.S. wrote and revised the manuscript. All authors reviewed and approved the manuscript. Dozens of participants of the Quartet Project freely donated their time and reagents for the completion and analysis of the project.

## Additional information

## Extended Data Figure Legends

**Extended Data Fig. 1.**
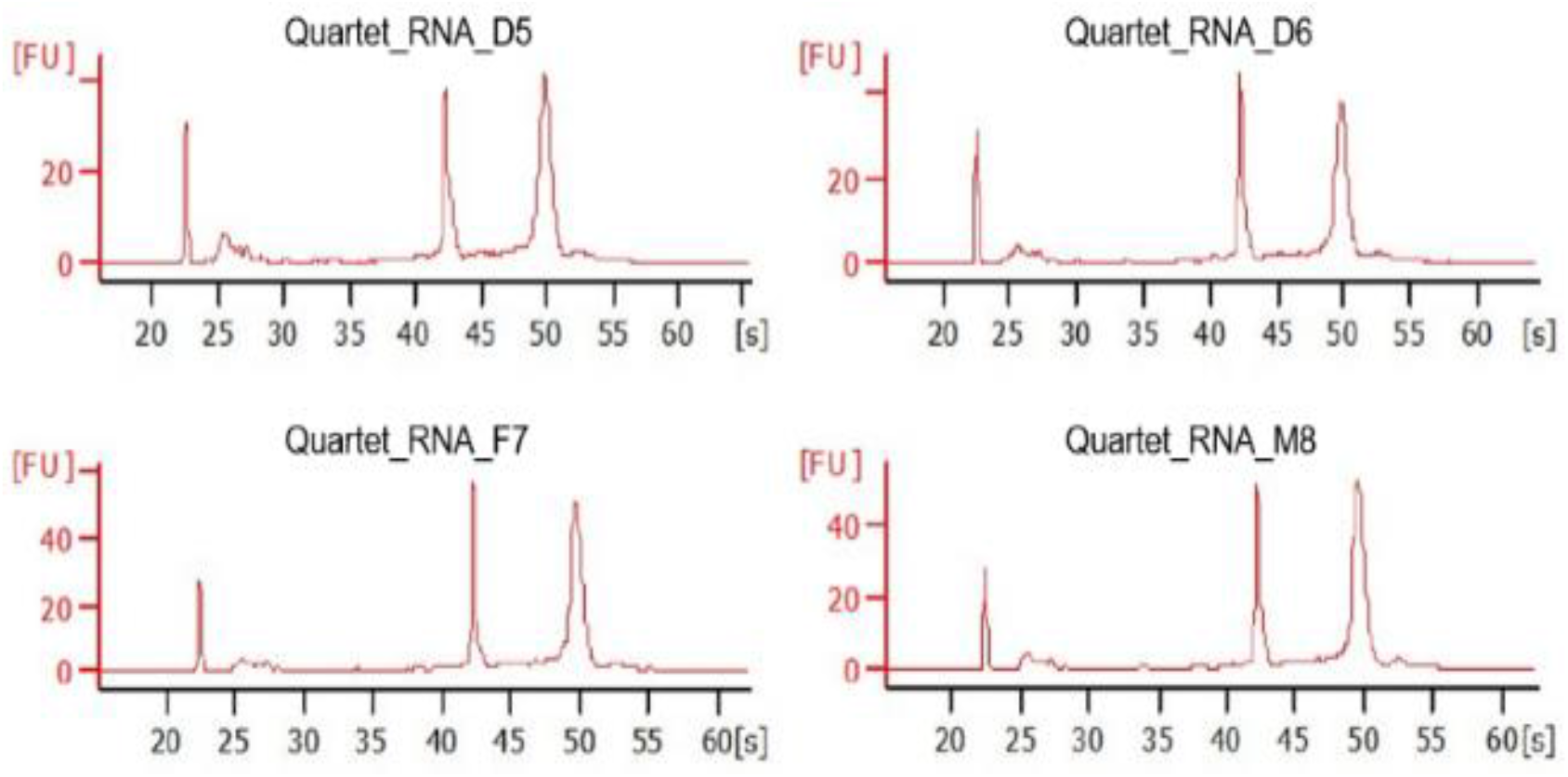
Quality of the Quartet RNA reference materials. Electropherograms of the Quartet RNA reference materials reflecting RNA quality produced by Agilent 2100.

**Extended Data Fig. 2.**
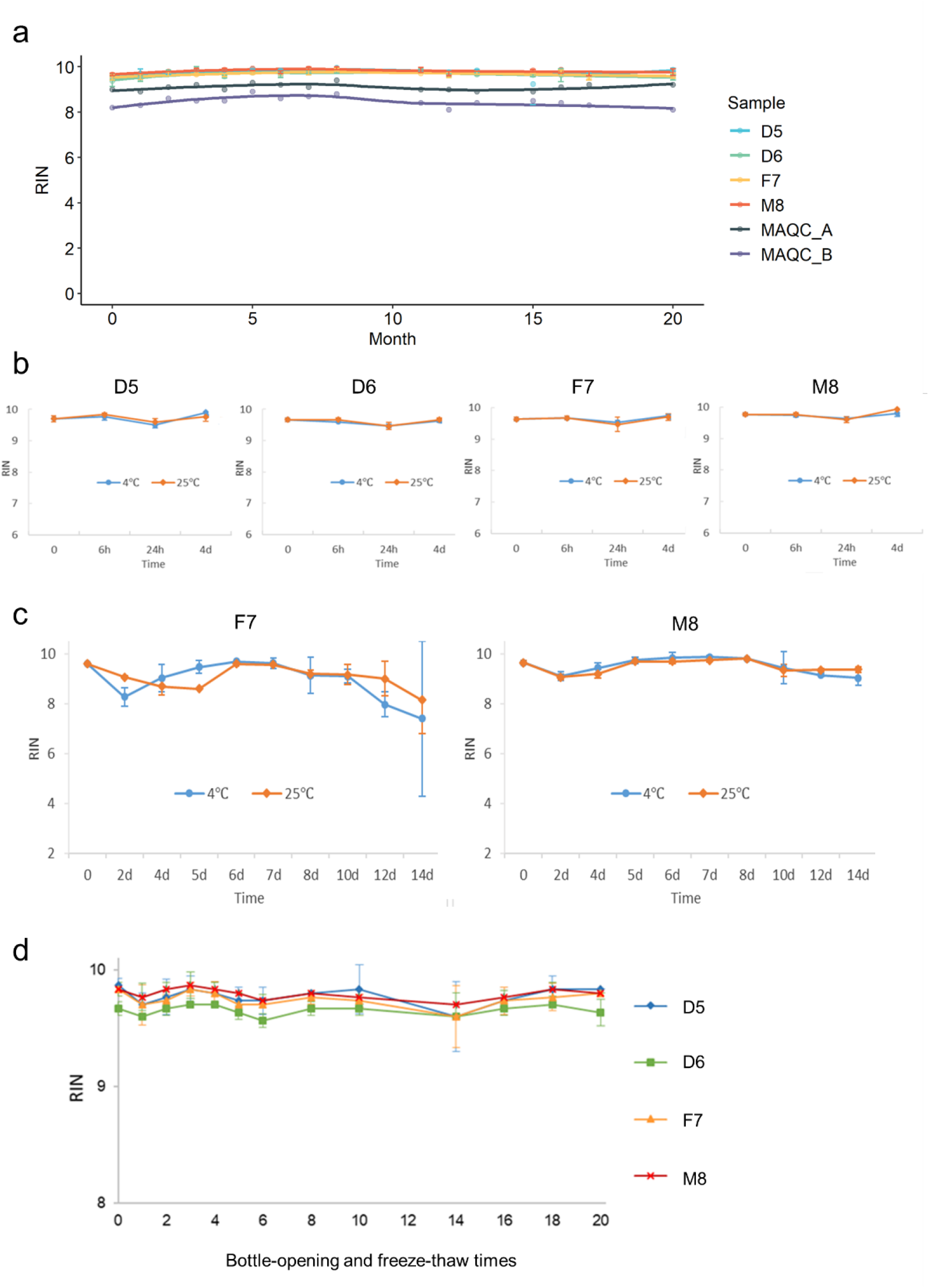
Stability assessment of the Quartet RNA reference materials. (**a**) Distribution of RNA integrity number (RIN) values across 20 months during assessment. The Quartet RNA reference materials and two well-characterized and commercialized RNA reference materials (MAQC A and B) were tested. (**b**) RIN values of the Quartet RNA reference materials across four days of storage at 4°C or 25°C. (**c**) RIN values of two reference materials (F7 and M8) across 14 days of storage at 4°C or 25°C. (**d**) RIN values of 20 times of bottle-opening and freeze-thaw cycle.

**Extended Data Fig. 3.**
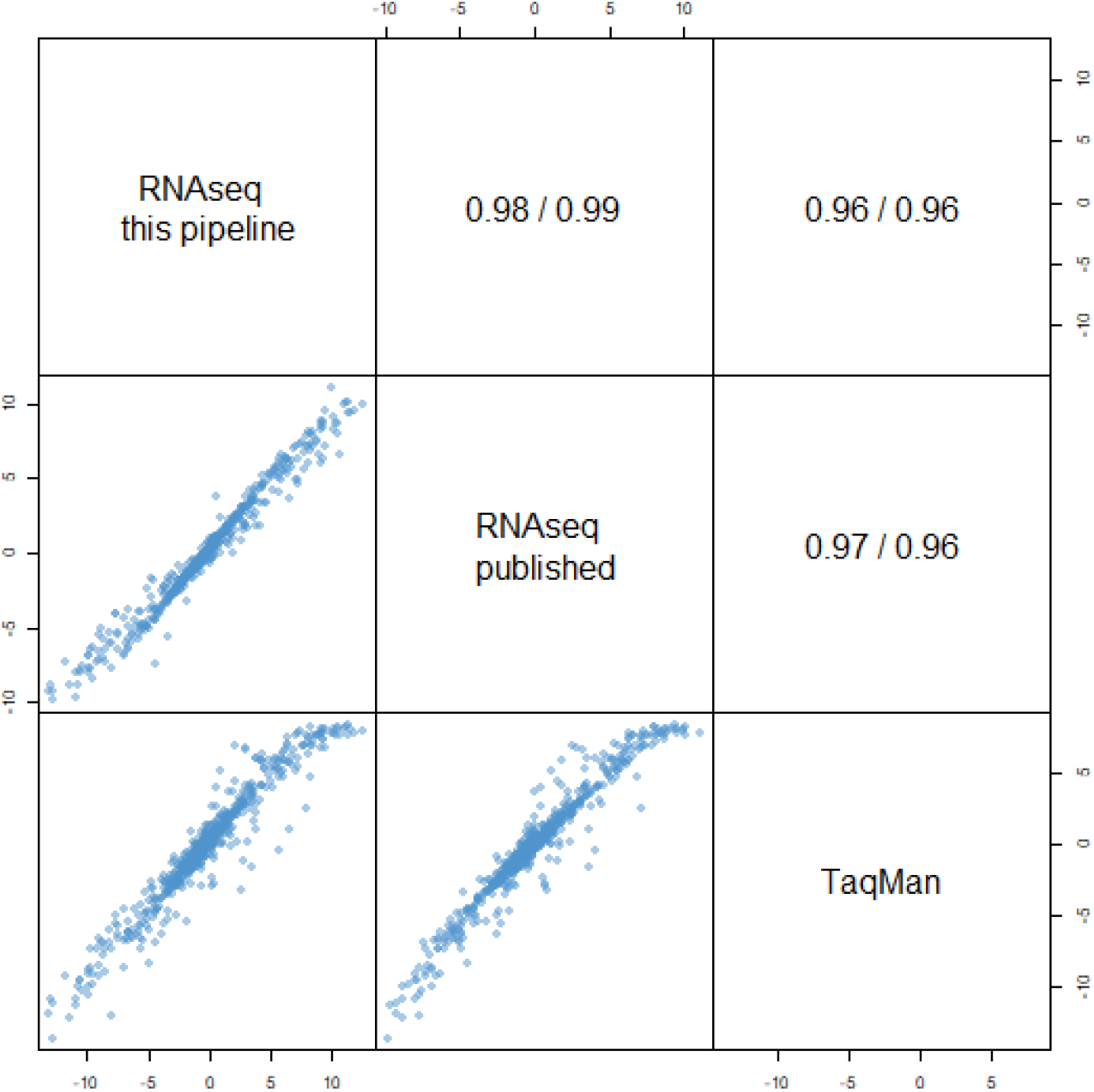
Validation analysis pipeline based on MAQC reference materials. The scatter plots compared the log2 fold differences or ratios (using MAQC B/A replicates) from RNAseq results using analysis pipeline used in this study, RNAseq results from publication^18^, and expression profiles obtained by MAQC-I TaqMan assays. Raw fastq files from public data (GSE47774) were used. Good concordances were observed between RNAseq and MAQC-I TaqMan assays^13^. The results revealed that analysis pipelines in this study were reliable. Pearson/Spearman correlation coefficients were shown in the upper-right triangle.

**Extended Data Fig. 4.**
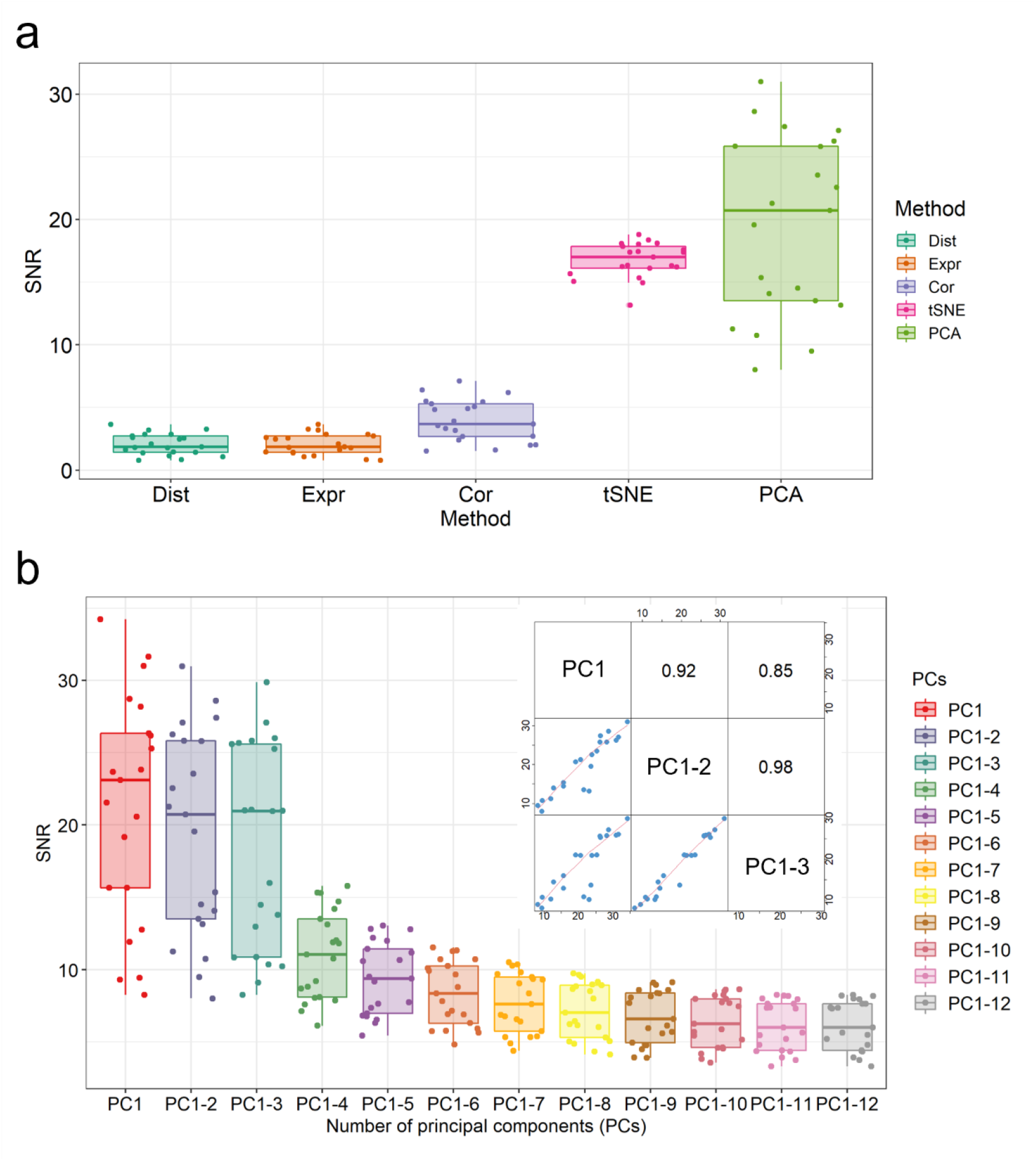
SNR is established for quality assessment across multiple samples. (**a**) Boxplots of distribution of SNR values calculated by different distance metrics, including Euclidean distance (Dist), overall expression profiles (Expr), Pearson correlation coefficient (Cor), t-Distributed Stochastic Neighbor Embedding (tSNE), and principal components analysis (PCA). (**b**) Boxplots of distribution of SNR values across 21 RNAseq batches calculated by using difference numbers of principal components.

**Extended Data Fig. 5.**
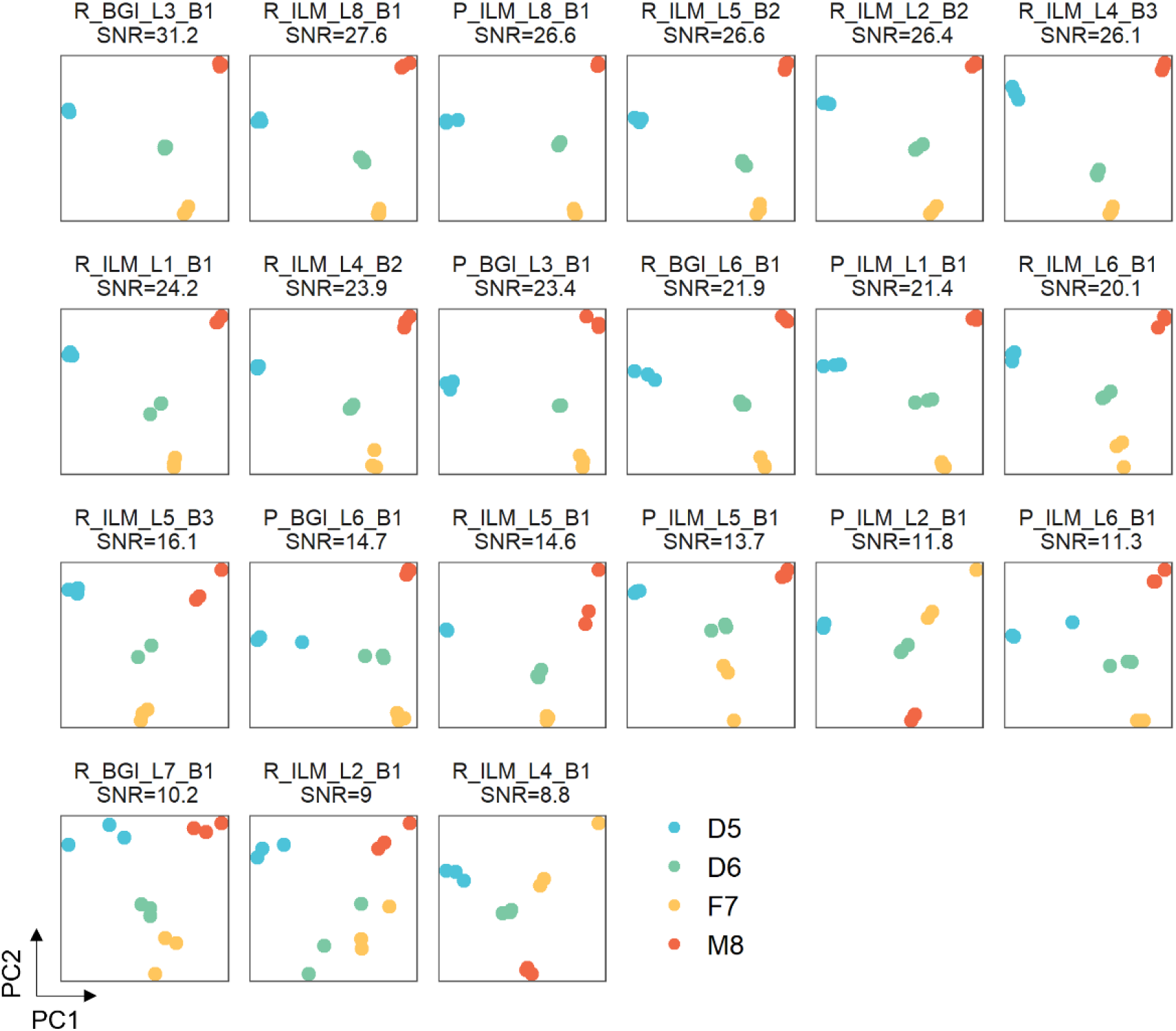
Intra-batch PCA plots across 21 RNAseq batches. Scatter plots of principal component analysis (PCA) in each batch based on log_2_FPKM normalized gene expression data. PCA plots were ordered by signal-to-noise ratio (SNR) values. Plots were color-coded by sample groups.

**Extended Data Fig. 6.**
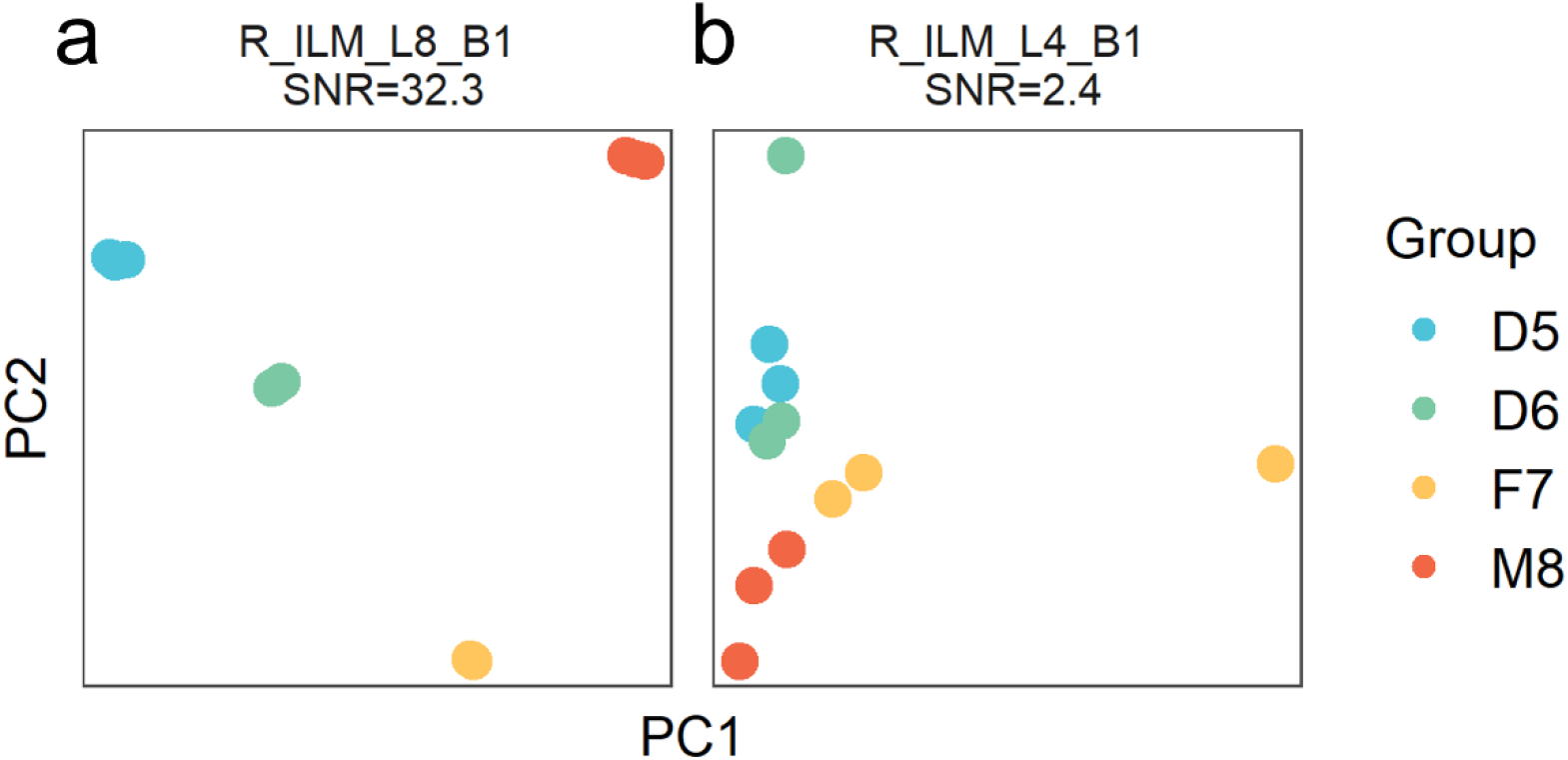
PCA of two exemplary batches based on Percent Spliced In (PSI) values of alternative splicing events. RNAseq datasets from (**a**) R_ILM_L8_B1 (as a high-quality batch) and (**b**) R_ILM_L4_B1 (as a low-quality batch) were used for plotting.

**Extended Data Fig. 7.**
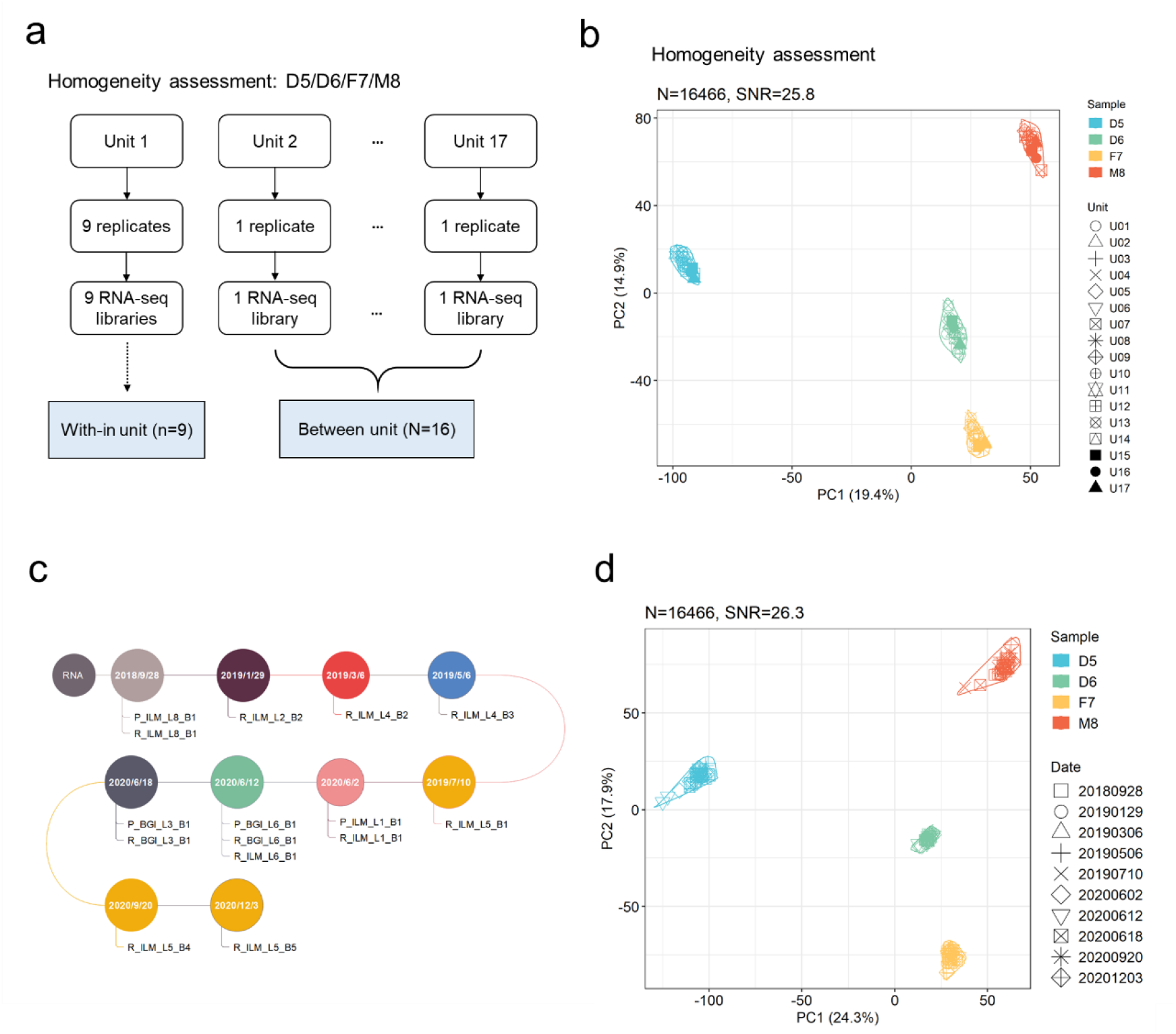
Assessment of homogeneity and long-term stability of the Quartet RNA reference materials. (**a, c**) Schematic diagrams of assessing homogeneity (**a**) and long-term stability (**c**) using RNAseq data. (**b**) Scatter plot of principal component analysis (PCA) of expression profiles across 17 units for homogeneity assessment. Plot was color-coded by sample groups and shaped-coded by packaging ID. (**d**) Scatter plot of PCA of expression profiles from 15 batches of data generated from up to 26 months. Plot was color-coded by sample groups and shaped-coded by date of data generation. Ratio-based expressions were obtained by subtracting log_2_FPKM by the mean of log_2_FPKM of the three replicates of D6 in the same batch.

**Extended Data Fig. 8.**
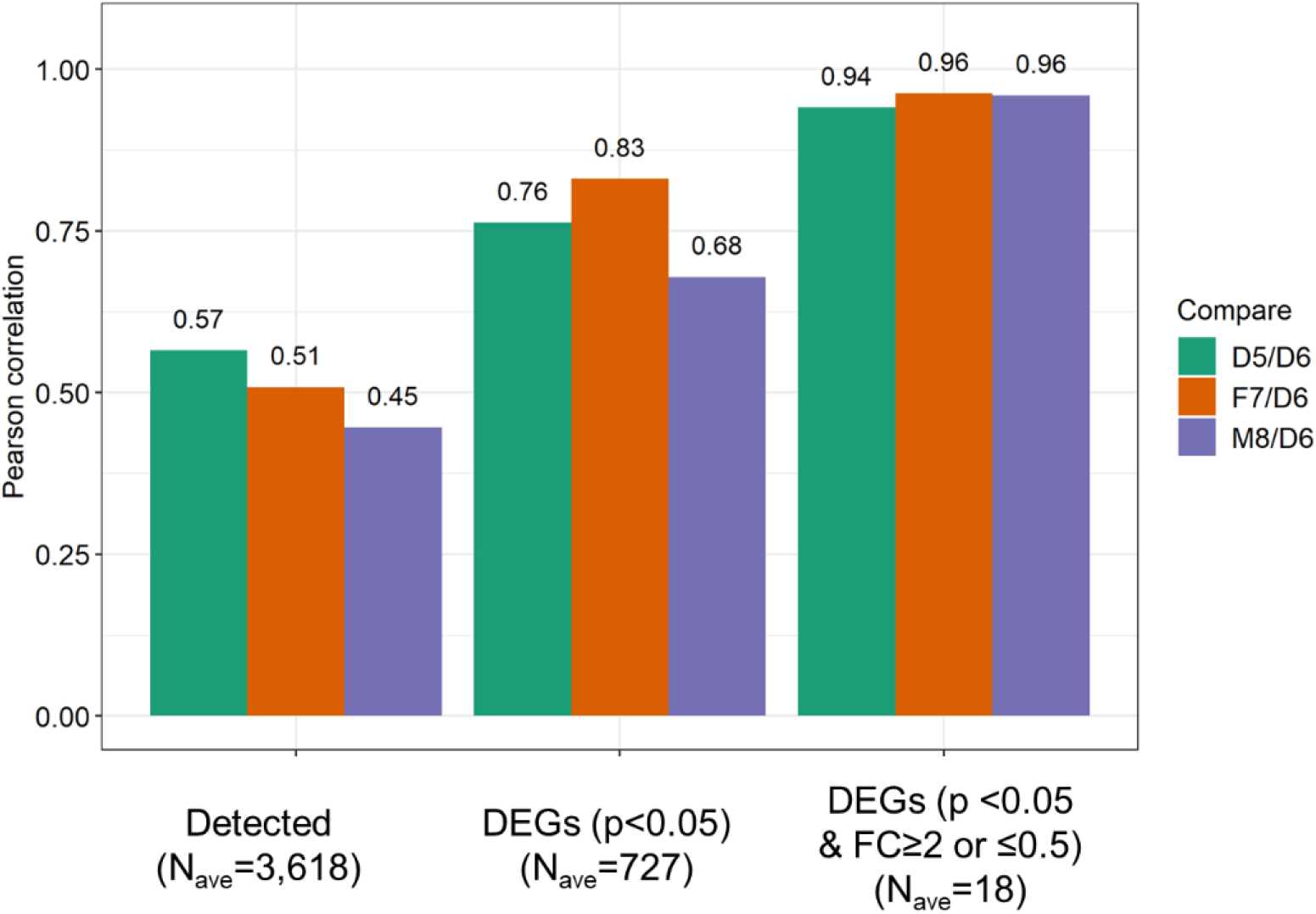
Correlation of RNA-protein pairs under different criteria for selecting the number of features for plotting. A batch of LC-MS/MS proteomic dataset was used for cross-omics validation of RNA reference datasets. Pearson correlation coefficient was used to represent consistency between RNA and protein abundances. RNA-protein correlations varied considerably when different cut-offs were used to select genes for plotting. Three different cutoffs were used with increasing degree of stringency, including detectable genes, differentially expressed genes based on *p*-value <0.05, or differentially expressed genes based on *p*-value <0.05 and fold change ≥ 2 or ≤ 0.5. The average numbers of genes across the three sample pairs were listed in the X-axis.

**Extended Data Fig. 9.**
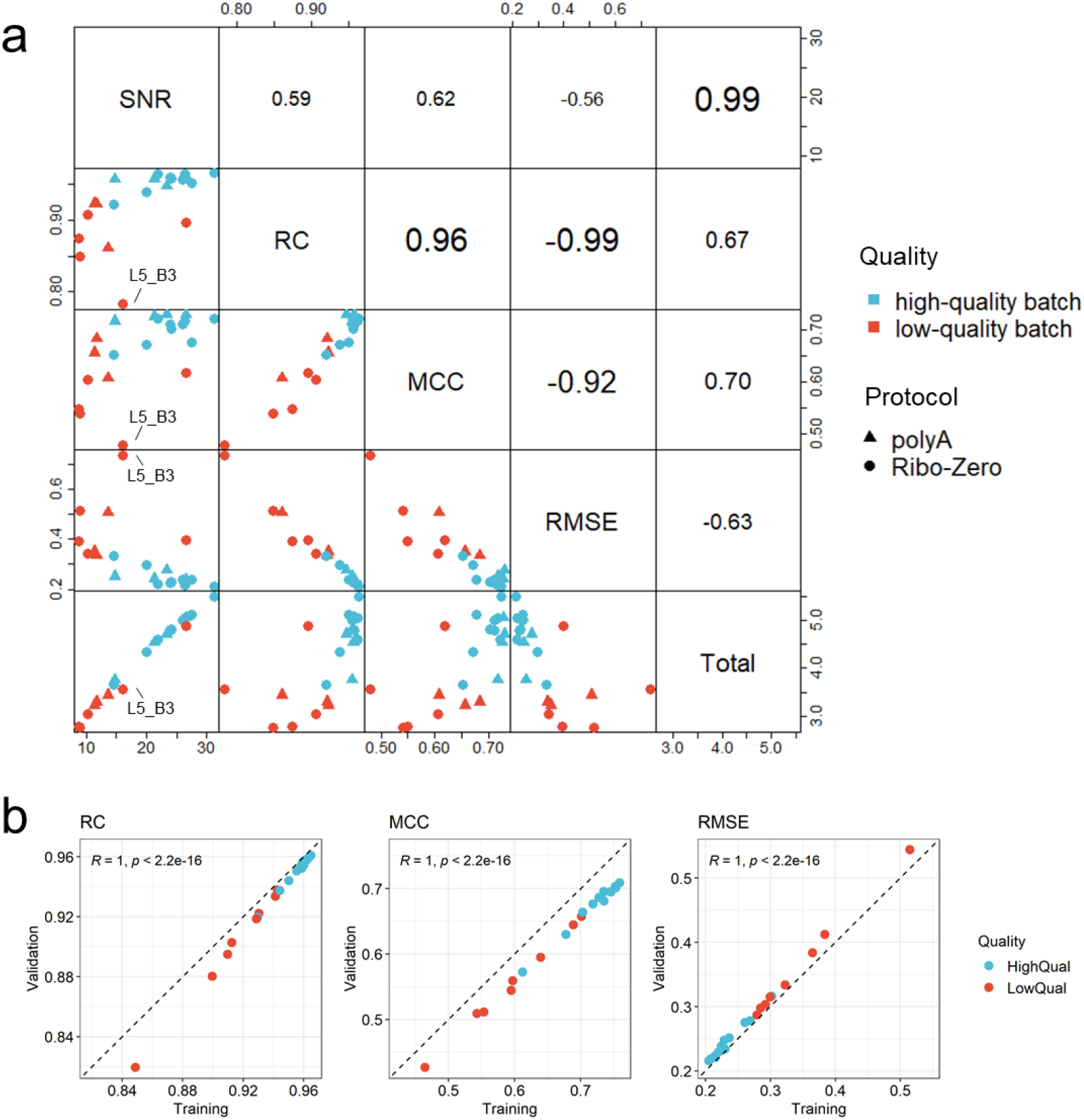
Concordance between different quality control metrics based on reference datasets. **(a)** Distribution of relative correlation with reference datasets (RC), MCC of DEGs, and root mean square error (RMSE) of the differences with reference datasets across 21 RNAseq batches. (**b**) Cross-validation test to estimate and validate the suitability of quality metrics with RC, RMSE and MCC of DEGs.

**Extended Data Fig. 10.**
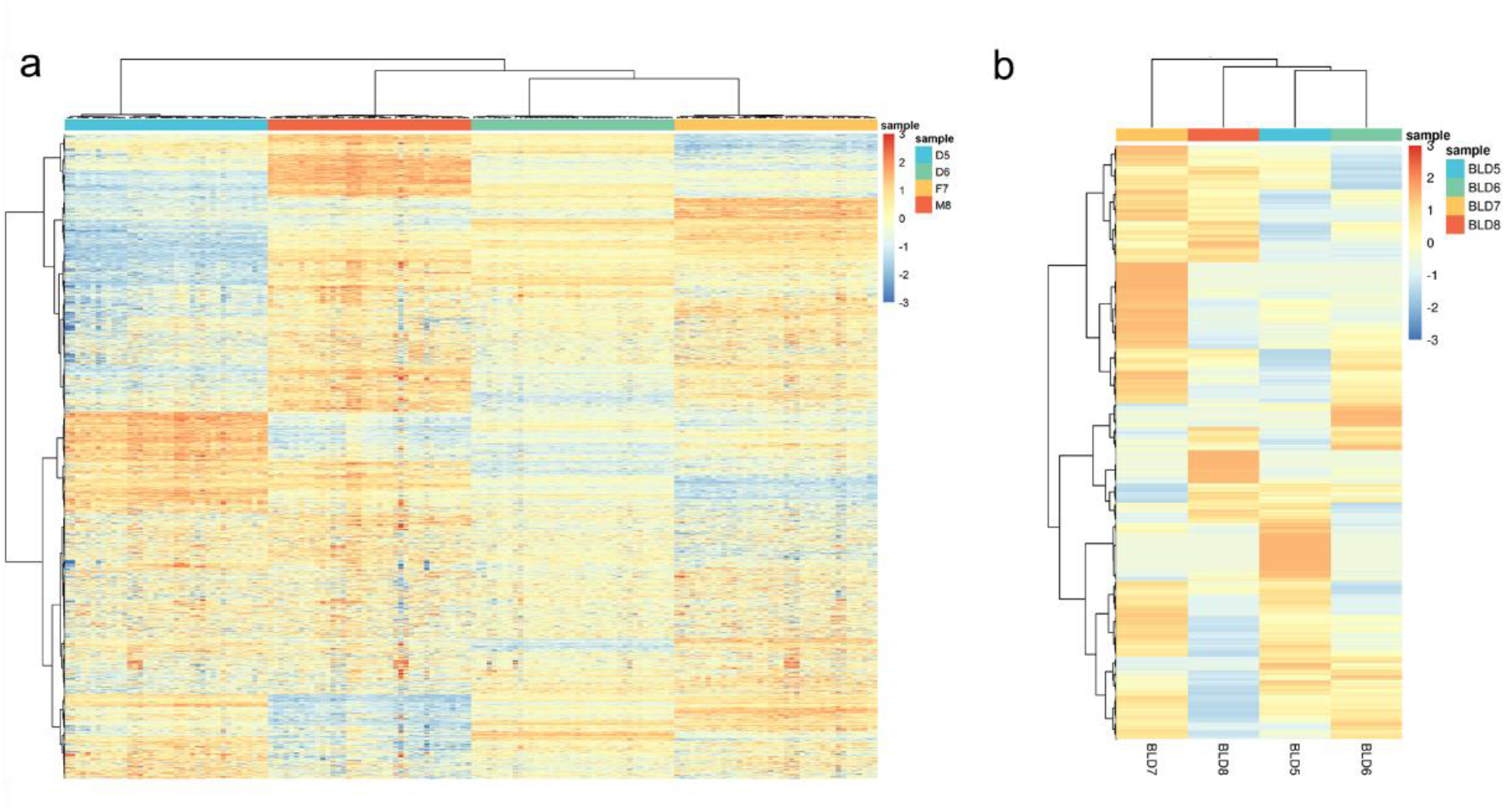
Hierarchical clustering of the immortalized cell lines and corresponding whole-blood samples of the quartet family members. **(a)** Hierarchical clustering based on 156 RNAseq libraries from 13 batches with high quality derived from the four immortalized cell lines of the quartet family members. Ratio-based expressions of detected genes across all samples were used (N=19,760). **(b)** Hierarchical clustering based on transcriptomic profiles of four whole-blood samples from the quartet family members. The overall expression profiles of detected genes (N=39,004) were used.

## Supplementary information

**Supplementary Table 1. Quality assessment of the Quartet RNA reference materials**

**Supplementary Table 2. Metadata of the Quartet RNAseq datasets**

**Supplementary Table 3. Quality control measurements across RNAseq batches**

**Supplementary Table 4. Ratio-based expressions of reference datasets and their extended uncertainties**

**Supplementary Table 5. Ratio-based expressions of reference datasets across 13 high-quality batches**

**Supplementary Table 6. Homogeneity, stability, and uncertainty assessment of the reference datasets**

**Supplementary Table 7. List of reference differentially expressed genes (DEGs) and non-differentially expressed genes**

**Supplementary Table 8. Genes and their primers used for qPCR validation**

**Supplementary Table 9. Fold changes and *p*-values of DEGs tested in reference datasets and qPCR results**

